# Cellular Modeling of CLN6 with IPSC-derived Neurons and Glia

**DOI:** 10.1101/2024.01.29.577876

**Authors:** Maria Gabriela Otero, Jaemin Kim, Yogesh Kumar Kushwaha, Alex Rajewski, Fabian David Nonis, Chintda Santiskulvong, Serguei I. Bannykh, Hiral Oza, Hafiz Muhammad Umer Farooqi, Madeline Babros, Christina Freeman, Lucie Dupuis, Saadat Mercimek-Andrews, Roberto Mendoza-Londono, Catherine Bresee, David R. Adams, Cynthia J. Tifft, Camilo Toro, Negar Khanlou, William A. Gahl, Noriko Salamon, Tyler Mark Pierson

## Abstract

Neuronal ceroid lipofuscinosis (NCL), type 6 (CLN6) is a neurodegenerative disorder associated with progressive neurodegeneration leading to dementia, seizures, and retinopathy. *CLN6* encodes a resident-ER protein involved in trafficking lysosomal proteins to the Golgi. CLN6p deficiency results in lysosomal dysfunction and deposition of storage material comprised of Nile Red^+^ lipids/proteolipids that include subunit C of the mitochondrial ATP synthase (SUBC). White matter involvement has been recently noted in several CLN6 animal models and several CLN6 subjects had neuroimaging was consistent with leukodystrophy. CLN6 patient-derived induced pluripotent stem cells (IPSCs) were generated from several of these subjects. IPSCs were differentiated into oligodendroglia or neurons using well-established small-molecule protocols. A doxycycline-inducible transgenic system expressing neurogenin-2 (the I3N-system) was also used to generate clonal IPSC-lines (I3N-IPSCs) that could be rapidly differentiated into neurons (I3N-neurons). All CLN6 IPSC-derived neural cell lines developed significant storage material, CLN6-I3N-neuron lines revealed significant Nile Red^+^ and SUBC^+^ storage within three and seven days of neuronal induction, respectively. CLN6-I3N-neurons had decreased tripeptidyl peptidase-1 activity, increased Golgi area, along with increased LAMP1^+^ in cell bodies and neurites. SUBC^+^ signal co-localized with LAMP1^+^ signal. Bulk-transcriptomic evaluation of control- and CLN6-I3N-neurons identified >1300 differentially-expressed genes (DEGs) with Gene Ontogeny (GO) Enrichment and Canonical Pathway Analyses having significant changes in lysosomal, axonal, synaptic, and neuronal-apoptotic gene pathways. These findings indicate that CLN6-IPSCs and CLN6-I3N-IPSCs are appropriate cellular models for this disorder. These I3N-neuron models may be particularly valuable for developing therapeutic interventions with high-throughput drug screening assays and/or gene therapy.

## 1. Introduction

Neuronal ceroid lipofuscinoses (NCLs) are the most common form of inherited pediatric neurodegenerative disease (Haltia and Goebel, 2013; Tuermer et al., 2021). Collectively termed Batten Disease, NCLs are 13 heterogeneous neurogenetic disorders associated with similar clinical and pathological features. Clinical features include progressive cognitive and motor decline, seizures, blindness, and an early death (Cárcel-Trullols et al., 2015; Haltia and Goebel, 2013), while pathological features include intracellular accumulation of lipid/proteolipid storage material. The genes/proteins involved in NCLs intersect with their involvement in the endolysosomal system and include catabolic lysosomal acid hydrolases (LAHs; CLN1, CLN2, CLN10, CLN13) along with several membrane-bound proteins involved in trafficking or other non-catabolic functions (CLN5, CLN6, CLN8)(Rechtzigel et al., 2022). Neuronal ceroid lipofuscinosis, type 6 (CLN6) is caused by bi-allelic pathogenic variants in the *CLN6* gene, which encodes a ubiquitously-expressed resident endoplasmic reticulum (ER) protein (CLN6p, Uniprot # Q9NWW5), which is involved in trafficking lysosomal proteins from the ER to the Golgi and eventual delivery to lysosomes (Bajaj et al., 2020; Gao et al., 2002).

Clinically, CLN6-related phenotypes are categorized into two major subtypes: a severe childhood form, CLN6A (or variant late-infantile (vLINCL); OMIM 601780)(Cárcel-Trullols et al., 2015; Haltia and Goebel, 2013; Rus et al., 2022; Tuermer et al., 2021) and an adult-onset disease, CLN6B (or Kufs type; OMIM 204300). Currently, no effective therapeutic interventions have been developed for CLN6A. The CLN6A subtype typically presents with delayed language development, cognitive regression, and motor decline (Canafoglia et al., 2015; Rus et al., 2022). Previous neuroimaging studies of patients clinically-classified with vLINCL reported primary involvement of cortical and subcortical grey matter (Badilla-Porras et al., 2022; Baker et al., 2017; Cannelli et al., 2009; Guerreiro et al., 2013; Jadav et al., 2014; Petersen et al., 1996; Peña et al., 2001; Peña et al., 2004); however confirmed genetic diagnoses were not always established in these subjects, so the data could be representative of a mix of clinically-similar NCLs (Baker et al., 2017; Jadav et al., 2014). Recently, a few genetically-confirmed CLN6 cases were also reported to possess white matter signal abnormalities (WMSAs) (Guerreiro et al., 2013; Biswas et al., 2020; Cannelli et al., 2009; Sun et al., 2018). This is consistent with recent reports of WMSAs being present in several CLN6 animal models (mouse, cat and sheep) (Sawiak et al., 2015; Faller et al., 2015; Johnson et al., 2020; Katz et al., 2020; Thelen et al., 2012a; White et al., 2018), indicating that white matter involvement may be a common part of the human disorder that requires further documentation.

Although CLN6p is ubiquitously expressed, most cellular pathology is limited to the CNS. Cortical neurons (layers II-VI), the hippocampus, and Purkinje cells are especially affected (Palmer et al., 1989; Marotta et al., 2017; Mole et al., 2005; Thelen et al., 2012a; Thelen et al., 2012b) and correlates with the most significant accumulation of storage material (Thelen et al., 2012a). To date, the specific molecular mechanisms of the neuronal disease are unclear, although studies performed with liver cells indicated that CLN6p works in tandem with CLN8p to traffic lysosomal proteins from the ER to the Golgi apparatus (Bajaj et al., 2020). In this non-neuronal model, the absence of CLN6p is thought to obstruct the export of LAHs and other essential lysosomal proteins from the ER, causing a deficiency of LAHs that mimics other NCLs that are associated with the deficient LAHs (CLN1, CLN2, CLN10)(Bajaj et al., 2020).

Alternatively, this mistrafficking could also lead to increased ER stress and activation of the Unfolded Protein Response (UPR)(Bajaj et al., 2020; Lakpa et al., 2021; Lojewski et al., 2014; Rutkowski and Kaufman, 2004). Lysosomal and ER stress both activate TFEB/TFE3 to translocate to the nucleus and promote transcription of CLEAR Network genes involved in lysosomal biogenesis and the UPR (Lojewski et al., 2014; Martina et al., 2014; Sardiello et al., 2009). These TFEB/TFE3 responses are valuable in the early stages of cellular dysfunction to stimulate pathways that alleviate these stressors including increased lysosomal biogenesis and enhanced cell division (Napolitano and Ballabio, 2016); however, chronic stimulation of TFEB/TFE3 responses can then activate signals associated with apoptosis (Bajaj et al., 2020; Martina et al., 2016; Rutkowski and Kaufman, 2004). In neurons, disruption of these endosomal/vesicular processes can be particularly toxic as critical distal functions within axons and synapses require intact intracellular protein trafficking (Ahrens-Nicklas et al., 2022; Ahrens-Nicklas et al., 2019; Rechtzigel et al., 2022).

CLN6 lysosomal dysfunction leads to the accumulation of lipid, protein, and proteolipid storage material (Haltia, et al., 2003; Cho et al., 2015; Seehafer and Pearce, 2009; Sima et al., 2018). The major proteinaceous storage component is subunit C of mitochondrial ATP-synthase (SUBC), making it a well-established biomarker of pathology (Best et al., 2017; Cao et al., 2011; Cárcel-Trullols et al., 2015; Gao et al., 2002; Haltia and Goebel, 2013; Jolly et al., 1992; Morgan et al., 2013; Oswald et al., 2005; Palmer et al., 1992; Palmer et al., 1989; Tammen et al., 2006; Thelen et al., 2012a). SUBC immunocytochemistry (ICC) has been a reliable method to detect protein storage, while lipophilic stains like Nile Red have also been used an additional simple and direct assay of lipid and proteolipid storage (Greenspan et al., 1985; Seehafer and Pearce, 2006; Sima et al., 2018). In addition to these methods, electron microscopy has revealed characteristic ultrastructural patterns associated with CLN6 (Sharp et al., 1997; Thelen et al., 2012a; Thelen et al., 2012b). Even though storage material has been widely used as a biomarker of the active disease process, there is an ongoing debate about whether storage accumulation is the specific etiology of neuronal dysfunction or a more passive biomarker of the disease process. This is especially true of NCLs associated with proteins involved in protein trafficking where axonal and synaptic dysfunction is seen before storage accumulates (Ahrens-Nicklas et al., 2022; Ahrens-Nicklas et al., 2019; Gomez-Giro et al., 2019; Osório et al., 2009; Rechtzigel et al., 2022); nonetheless, storage biomarkers can be used to measure efficacy of any therapeutic interventions.

In this report, we describe new insights in both clinical and laboratory evaluations of CLN6A. We discuss our development of renewable patient-derived human induced pluripotent cell (IPSC) models of CLN6 and their utility to model disease pathology in neural cells. We also discuss the presence of white matter signal abnormalities (WMSAs) on neuroimaging of several genetically-confirmed CLN6A patients. Consistent with this finding, CLN6 IPSC-derived oligodendroglial cells were produced and found to possess increased SUBC+ storage material. We then used two different neuronal differentiation protocols to generate IPSC-derived neuronal cells. These methods included small molecule protocols (dual SMAD inhibition), as well as the I3N-system for inducible expression of neurogenin-2 (NGN2) for directed differentiation of IPSCs into neurons (I3N-neurons)(Zhang et al., 2013; Fernandopulle et al., 2018; Shi et al., 2012a; Shi et al., 2012b; Wang et al., 2017 and reviewed in Hulme et al., 2022). All CLN6-IPSC-derived neuronal cultures accumulated significant amounts of SUBC^+^ storage. Additional investigations with I3N-neurons revealed: i) ultrastructural characteristics present in other models and pathological samples of CLN6; ii) increased Nile Red^+^ lipid storage; iii) enlarged area of GM130^+^ (Golgi matrix protein 130) signal; iv) increased LAMP1^+^ (lysosomal associated membrane protein1) signal; and v) decreased tripeptidyl peptidase-1 (TPP1) activity. Bulk transcriptomic data analysis comparing CTL- and CLN6-I3N-neurons identified a large number of differentially-expressed genes (DEGs) involved in biological processes or pathways affecting neuronal apoptosis, intracellular transport, and axonal and synaptic organization/function as identified by Gene Ontology (GO) and Canonical Signaling pathway analysis.

## 2. Materials and methods

### 2.1 Patient recruitment and ethics approval

Patients were enrolled in protocols 76-HG-0238 and Pro00038462 approved by the National Human Genome Research Institute (NHGRI) Institutional Review Board and the Cedars-Sinai Medical Center Institutional Review Board, respectively.

### 2.2 IPSC Generation, Cell Culture techniques and Cell-Type Specific Differentiation

Blood samples were provided by four CLN6 and four CTL subjects enrolled in our study. IPSCs were generated from blood samples as per previously published protocols (Barrett et al., 2014). Pluripotency was validated by a high nuclear-to-cytoplasmic ratio, positivity for alkaline phosphatase and the surface antigens SSEA4, TRA-1-60, TRA-1-81, and the presence of nuclear pluripotency markers OCT3/4, SOX2 and NANOG. All the IPSC lines possessed a normal G-band karyotype. All IPSC lines strictly fulfilled the standard criteria of successful cellular reprogramming without vector integration as defined by the downregulation of exogenous reprograming factors and concurrent expression of factors that include pluripotency markers (NANOG, SSEA4, TRA1) and known IPSC markers (SOX2, NANOG, LIN28, GDF3, OCT4, DNMT3B) as determined by semiquantitative RT-PCR. Pluritest (ThermoFisher Scientific, Waltham, MA) was used to assign a novelty score in which all IPSC lines were highly dissimilar to the non-pluripotent samples, as well as a pluripotency score that demonstrated that all lines were pluripotent. Hierarchical clustering of the samples based on PluriTest Gene Expression profile shows that IPSCs clustered close to each other and away from human neural progenitor cells (NPCs) and fibroblasts. Southern blotting was used to verify that there was no integration of the plasmids used to generate the IPSCs (Barrett et al., 2014; Mattis et al., 2015). All lines were routinely tested for mycobacteria, infectious agents, and other potential contaminants.

Differentiation into IPSC-derived oligodendroglia: IPSCs were differentiated into oligodendroglial cells with a well-established protocols to evaluate for storage accumulation (Hu et al., 2009a; Hu et al., 2009b; Wang et al., 2013) CTL- and CLN6-IPSCs were grown under standard conditions until they became confluent. At that point all cleaned colonies were detached with Accutase (Stemcell Technologies, Cambridge, MA) and plated into a poly-hema treated 6-cm Petri dish in media containing mTeSR1 (StemCell Technologies, Cambridge, MA) and 10μM rock inhibitor (RI) (StemGent, Beltsville, MD). After 2 days, embryoid bodies (EBs) were collected in a 15-ml Falcon tube by gravity flow. Media was aspirated and collected EBs were resuspended in fresh Neuronal differentiation medium (NDM) containing DMEM/F12 (Gibco), 0.25% penicillin/streptomycin (Gibco), 1% N2 supplement (Gibco), 1% NEAA (Gibco), 2mg/ml Heparin (Sigma-Aldrich, St. Louis, MO), 1μM cAMP (Sigma-Aldrich, St. Louis, MO), 20ng/ml Ascorbic acid (Sigma-Aldrich, St. Louis, MO) and supplemented with 10ng/ml IGF-1 (Hu et al., 2009a; Hu et al., 2009b; Wang et al., 2013). Media was changed in this manner on day 4. On Day 6 of EB formation, collected EBs were resuspended in NDM and 10ng/ml IGF-1 and plated onto 0.5mg Matrigel-coated (Corning, Corning, NY) 6-well plates. The plates were incubated at 37°C and left undisturbed for 3 days for attachment. On day 14, neural rosette aggregates were picked using StemDiff Neural rosette selection reagent (Stemcell technologies, Cambridge, MA).

Collected rosettes were plated onto a polyhema-treated 6-cm Petri dish in NDMB medium containing NDM and 2% B27 w/o vitamin A (Gibco, Grand Island, NY), supplemented with 10ng/ml IGF-1, 0.1μM RA (Reprocell, Beltsville, MD), 1μM purmorphamine (Apexbio, Houston, TX) and 10μM RI (StemGent, Beltsville, MD). Media (without RI) was changed every 2 days, and neurospheres that were formed in suspension were disaggregated into smaller cell clusters by rapid pipetting. Differentiation cultures at Day O24 were treated with NDMB (supplemented with 10ng/ml IGF-1, 1μM Purmorphamine (Apexbio, Houston, TX) and 10ng/ml FGF2 (Peprotech, Westlake Village, CA)) (Hu et al., 2009a; Hu et al., 2009b; Wang et al., 2013). Only half of the media was changed every 2 days. At Day O32, Pre-Oligodendrocyte Progenitor Cells (Pre-OPCs) were differentiated to Oligodendrocyte Progenitor Cells (OPCs) by using Glial Differentiation Medium (GDM) that contained DMEM/F12 (Gibco, Grand Island, NY), 0.25% penicillin/ streptomycin (Gibco, Grand Island, NY), 2% B27 w/o vitamin A (Gibco, Grand Island, NY), 1% N1 supplement (Sigma-Aldrich, St. Louis, MO), 1% NEAA (Gibco, Grand Island, NY), 1μg/ml Biotin (ThermoFisher Scientific, Waltham, MA), 1μM cAMP (Sigma-Aldrich, St. Louis, MO), 60ng/ml T3 and supplemented with 10ng/ml IGF-1, 1μM purmorphamine (Apexbio, Houston, TX), 10ng/ml PDGF-AA (100-13A, PeproTech, Westlake Village, CA) and 10ng/ml NT3 (Peprotech, Westlake Village, CA)(Hu et al., 2009a; Hu et al., 2009b; Wang et al., 2013).

Media (without RI) was changed every 2 days and gliogenic spheres in suspension were disaggregated into smaller cell clusters by rapid pipetting. At Day O46, in order to differentiate OPCs into immature oligodendrocytes, the media was changed to GDM supplemented with 10ng/ml IGF-1, 10ng/ml PDGF-AA (Peprotech, Westlake Village, CA), 10ng/ml NT3 (Peprotech, Westlake Village, CA)(Hu et al., 2009a; Hu et al., 2009b; Wang et al., 2013), that was changed every 2 days. For final differentiation into Oligodendrocytes at Day O82, around 50 cell clusters were seeded onto 0.5mg Matrigel-coated 6-well plates provided with fresh GDM supplemented with 10ng/ml IGF-1, 1μM purmorphamine (Apexbio, Houston, TX), 10ng/ml PDGF-AA (Peprotech, Westlake Village, CA) and 10ng/ml NT3 (Peprotech, Westlake Village, CA). The next day, the media was changed to GDM supplemented with 5ng/ml IGF-1, 1μM purmorphamine (Apexbio, Houston, TX), 5ng/ml PDGF-AA (Peprotech, Westlake Village, CA) and 5ng/ml NT3 (Peprotech, Westlake Village, CA), which was changed every 2 days until reaching Day O110 (Hu et al., 2009a; Hu et al., 2009b; Wang et al., 2013).

Cultures ready for harvest were dissociated two days before each assigned timepoint (Day O32, O46, O82 and O110) to be plated as monolayers (Hu et al., 2009a; Hu et al., 2009b; Wang et al., 2013). Cell clusters floating in suspension were dissociated into single-cells with Accutase (Stemcell technologies, Cambridge, MA). The single cell population was resuspended in their respective NDM/GDM media supplemented with RI (StemGent, Beltsville, MD). The resuspended cells were then seeded onto poly-L-ornithine/ Matrigel-coated coverslips in 24-well plates at a density of 50,000-100,000 cells/well for two days and then underwent ICC evaluations (see Section 2.3) (Hu et al., 2009a; Hu et al., 2009b; Wang et al., 2013).

Differentiation into IPSC-derived neuronal cultures using dual SMAD inhibition: IPSCs were differentiated using previously published protocols with some modifications (Shi et al. 2012). Briefly, CTL- and CLN6-cells were grown under standard conditions until they became ∼80% confluent. Cells then were detached using Dispase (StemCell, Cambridge, MA) and plated again in a dilution 1:6 in neural differentiation medium to induce neuroepithelial differentiation (NDM; 50% DMEM-F12 (Gibco, Grand Island, NY), 50% Neurobasal (Gibco, Grand Island, NY), 2.5 μg/ml human insulin (Sigma-Aldrich, St. Louis, MO), 100 μM ß-mercaptoethanol (Sigma-Aldrich, St. Louis, MO), 1x non-essential amino acid (NEAA; Gibco, Grand Island, NY), 0.25x penicillin/streptomycin (Gibco, Grand Island, NY), 1x N2 (Life Technologies, Carlsbad, CA) and 1x B27 (Gibco, Grand Island, NY)) supplemented with 0.5 μM LDN193189 (Sigma-Aldrich,

St. Louis, MO) and 10 μM SB431542 (Tocris, Minneapolis, MN). Media was changed daily. At Day 7, the cells were passaged with Versene (ThermoFisher Scientific, Waltham, MA), and transferred onto fresh Matrigel (Corning, Corning, NY)-coated 6-well dishes. The following day, the medium was changed to NDM supplemented with 20 ng/ml fibroblast growth factor-2 (FGF2; Peprotech, Westlake Village, CA). Media was changed daily. After 4-6 days, the induced neural rosettes were isolated by a Neural Rosette Selection Reagent (Stem Cell Technologies, Cambridge, MA) and the isolated neural rosettes were transferred onto a fresh Matrigel-coated 6-well dish and expanded in NDM supplemented with FGF2 (Peprotech, Westlake Village, CA) for 4-6 days. The expanded neural rosettes were harvested by Accutase (Stem Cell Technologies, Cambridge, MA) and then frozen at −80°C for short-term storage or in a liquid nitrogen tank. The frozen neural rosettes were thawed and cultured in NDM supplemented with 10 μM RI (StemGent, Beltsville, MD) onto a fresh Matrigel-coated 6-well dish for 2-4 days to recover. The cells were harvested using Accutase (Stemcell Technologies, Cambridge, MA), resuspended in NDM supplemented with 10 μM RI (StemGent, Beltsville, MD), and seeded onto 4 fresh Matrigel-coated coverslips in a well of a 6-well dish at a density of 300,000 cells/well. This last step was performed on a day that was designated “Day Post-Thaw 1” or “Day PT1”, with subsequent days designated in order (Day PT2, Day PT3, etc.). Half of the media was changed every 2 days without RI. Neuronal cultures were harvested on representative timepoints (PT7, PT14, PT21, and PT28) and underwent ICC evaluations (see Section 2.3).

Generation of I3N-IPSCs and differentiation into I3N-neurons: The I3N-system was used to generate DOX-inducible IPSCs that would differentiate into I3N-neurons within 2-4 days of DOX-exposure (Fernandopulle et al., 2018; Wang et al., 2017). I3N-IPSC lines were generated from undifferentiated human CTL-(003, 201, 276 and 888) and CLN6-(121, 313, 770 and 776) IPSC lines per previously published protocols with some modifications (Fernandopulle et al., 2018; Wang et al., 2017). To summarize, undifferentiated IPSCs were cultured on Matrigel-coated dishes in mTeSR1 medium (StemCell Technologies, Cambridge, MA) and passaged as aggregates. In order to generate clonal I3N-IPSC lines, IPSC cells were dissociated as single cells with Accutase (Stemcell technologies, Cambridge, MA) and were transfected with plasmids pTALdNC-AAVS1_T2 (Addgene # 80496), pTALdNC-AAVS1_T1 (Addgene # 80495), and pUCM-AAVS1-TO-hNGN2 (Addgene #105840) in Lipofectamine stem transfection reagent (Life Technologies, Carlsbad, CA) and then plated fresh on Matrigel-coated 6-well dishes (1,500,000 cells/well). Transfected IPSCs were evaluated with fluorescence microscopy several days later to visualize transgene-containing mCherry^+^ cells. mCherry^+^ IPSCs were then manually isolated and expanded over several cycles to increase the percentage of mCherry^+^ IPSCs in culture before finite isolation by fluorescent cell activated sorting (FACS) using Influx cell sorter (BD Biosciences, San Diego, CA) with subsequent plating and expansion. Single colonies of mCherry^+^ cells were isolated and expanded and then analyzed for: i) neural differentiation by DOX treatment, ii) correct transgene integration into the adeno-associated virus integration site 1 (AAVS1) locus with PCR to identify appropriate clones, and iii) a normal karyotype (Fernandopulle et al., 2018; Wang et al., 2017). IPSCs that met all these requirements were then expanded and used for subsequent DOX-inducible differentiation and analysis. For DOX-inducible differentiation, the generated CLN6- and CTL-I3N-IPSCs were dissociated into single cells using Accutase (Stemcell technologies, Cambridge, MA) and resuspended in DOX-induction medium (DMEM-F12 (Gibco, Grand Island, NY) supplemented with 1% NEAA (Gibco, Grand Island, NY), 1% N2 (Gibco, Grand Island, NY), 2% B27 (Gibco, Grand Island, NY), 10 ng/ml brain-derived neurotrophic factor (BDNF; Prospect, East Brunswick NJ), 10 ng/ml neurotrophin-3 (NT3; Peprotech, Westlake Village, CA) and 2 μg/ml DOX (Sigma-Aldrich,

St. Louis, MO) and supplemented with 10 μM RI (StemGent, Beltsville, MD). Cells were then seeded at 400,000 cells per well of fresh Matrigel-coated 6-well dishes. The media was changed daily for 3 days without RI (Fernandopulle et al., 2018; Wang et al., 2017). For further maturation of induced I3N-neurons, the cells were dissociated with Accutase (Stemcell technologies, Cambridge, MA) and transferred onto fresh Matrigel-coated coverslip containing wells of a 24-well dish at a density of 50,000 cells/well in BrainPhys neuronal maturation medium (Stemcell Technologies, Cambridge, MA) supplemented with 1% N2 (Gibco, Grand Island, NY), 1% B27 (Gibco, Grand Island, NY), 10 ng/ml BDNF and 10ng/ml NT3 (Peprotech, Westlake Village, CA) supplemented with 10 μM RI (StemGent, Beltsville, MD)(Fernandopulle et al., 2018; Wang et al., 2017). This day was considered Day N1. Half of the media (without RI) was changed every 3-4 days (StemGent, Beltsville, MD). Cells were subsequently fixed at appropriate time points (N1, N3, N7, N14, and N21; see Section 2.3) for each associated experiment and evaluated with ICC, Nile Red staining, protein analysis, biochemical testing, or RNA analysis.

### 2.3 Immunocytochemistry and staining of samples

Coverslips were fixed in 4% paraformaldehyde (PFA; Electron Microscopy Sciences, Hatfield PA) at room temperature for 20 min, washed 3 times with PBS and permeabilized with 0.2%-Triton-100 in PBS for 10 min at room temperature, blocked with 5% goat serum (Gibco, Grand Island, NY) with 0.2%Triton-100 in PBS (Corning, Corning NY) for 1 hour at room temperature and then incubated overnight at 4°C with primary antibody. After 3 washes with PBS (Corning, Corning NY) coverslips were incubated with secondary antibody 2 hours at room temperature, washed 3 more times and mounted in Fluoromount-G (Southern Biotech, Birmingham, AL).

ICC of IPSC-derived Oligodendroglial cultures: CLN6- and CTL-IPSCs were differentiated into oligodendroglia as per above Section 2.2 (Hu et al., 2009a; Hu et al., 2009b; Wang et al., 2013). Cells were collected at representative time-points as follows: i) Pre-OPC stage (timepoint: Day O34) ii) OPC stage (timepoint: Day O48), iii) Immature Oligodendrocyte stage (timepoint: Day O84) and iv) Oligodendrocyte stage (timepoint: Day O112) (see Supplemental Fig 2). ICC testing was performed for SUBC at all timepoints and several oligodendroglial cell-type specific markers that included: Days O34 and O48: OLIG2 (pre-OPC marker) and NKX2.2 (OPC marker); Days O34, O48, O84, and O112: NG2 and PDGFRa (immature oligodendrocyte markers); and Days O84 and O112: O4 (oligodendrocyte marker).

ICC of IPSC-derived neuronal and I3N-neuronal cultures: CLN6- and CTL-IPSCs were differentiated into neuronal cultures as per above Section 2.2 (Shi et al. 2012). Cells were collected at representative time-points PT7, PT14, PT21 and PT28. ICC testing was performed for SUBC at all timepoints and neuronal markers (NeuN, TUJ1 or MAP2) at each timepoint.

CLN6- and CTL-I3N-IPSCs were differentiated into I3N-neurons as per above Section 2.2 (Fernandopulle et al., 2018; Wang et al., 2017). Cells were collected at representative time-points N1, N3, N5, N7, N14, and N21. ICC testing was performed for SUBC at all timepoints and neuronal markers (TUJ1 or MAP2) at each timepoint.

Primary antibodies: OLIG2 (1:100, Santa Cruz Biotechnology, Dallas, TX), NKX2.2 (1:200, DSHB, Iowa City IA), NG2/Anti-Chondroitin Sulfate (1:400, BD Biosciences, San Diego, CA), PDGFRα (1:50, Santa Cruz Biotechnology, Dallas, TX), Oligodendrocyte Marker O4 (1:100, R&D systems, Minneapolis, MN), SUBC (1:200, Abcam, Waltham, MA), Nestin clone 10C2 (1:1000, Millipore, Burlington, MA), B3-Tubulin (1:500; Millipore, Burlington, MA), MAP2ab (1:1000; Sigma-Aldrich, St. Louis, MO), LAMP1 (1:400; Santa Cruz Biotechnology, Dallas, TX), p62 (1:300 P62/SQSTM1, Sigma-Aldrich, St. Louis, MO), GM130 (1:300 BD Biosciences, San Diego, CA). Secondary antibodies: Alexa Fluor conjugated secondary antibodies (anti-mouse Alexa Fluor 488 and 568, anti-rabbit Alexa Fluor 488 and 568; ThermoFisher Scientific, Waltham, MA).

Nile Red staining was done per established protocols (Sima et al., 18) at appropriate timepoints. Nile Red stock solution (10ug/ml or 1ug/ml) was added to the appropriate media or PBS at a ratio of 1:1500 to make Nile Red Staining Solution. Media was then removed from tested cells and the Nile Red Staining Solution was added (for example 0.5ml for a 24 well plate well) and incubated at 37 degrees in incubator. After 15 minutes, remove the Nile Red Staining Solution and then fix the cells as per above. Samples stained with Nile Red could be directly visualized with microscopy or could also undergo ICC staining as per above as needed.

Where appropriate, samples were also counterstained with DAPI (4′,6-diamidino-2-phenylindole; ThermoFisher Scientific, Waltham, MA) to visualize nuclei and/or CellMask (ThermoFisher Scientific, Waltham, MA) to visualize the plasma membranes of cells as per product protocols. CellMask plasma membrane stains can deliver uniform staining of plasma membrane across a wide variety of mammalian cell types that are slow to internalize.

Slides and sections were imaged on a Nikon A1R laser-scanning confocal microscope with a 20x or 60x objective. ‘No antibody’ and ‘secondary-only’ controls were included in these experiments, and imaging settings were optimized to ensure autofluorescence and non-specific signals from secondary antibodies did not contribute to the immunofluorescence images. Maximum projecting fluorescence images were captured using the A1R confocal microscope with Nikon software (z-stack function with 8 stacks (2 μm apart from each other) at appropriate wavelengths for respective antibodies and dyes. Cells from four randomly chosen fields of vision were focused and captured. For quantifying SUBC puncta, GM130^+^ and LAMP1^+^ area, the captured images containing 70-100 cells per image were analyzed using ImageJ software (Grubman et al., 2014). Experiments from individual cell lines were run in at least triplicate samples. Statistical significance was determined by one-way ANOVA and Bonferroni multiple comparisons post hoc analysis using GraphPad Prism for SUBC analysis. For GM130 and LAMP1, longitudinal analysis of data across CTL- and CLN6-cell lines was performed with mixed model regression to account for the random effect of repeated measures from each individual cell line. Post-hoc pairwise testing for significant differences was Tukey corrected at the two-tailed alpha level of 0.05.

### 2.4 Electron microscopy

I3N-cells at Day N7 were washed twice with PBS and scraped off dish with a cell scrapper. The pellet collected was centrifuged 200g for one minute and fixed with 2.5% glutaraldehyde (GA) in sodium cacodylate buffer (SCB, ThermoFisher Scientific, Waltham, MA) for 30 min. Then the cells were scraped off, collected to 1.5 mL Eppendorf tubes, and serially pelleted in microcentrifuge at 13000 rpm over three-minute cycles with decanting of excess fixative and forming a tight pellet. Subsequently, the GA was washed off the pellet by serial overlay with SCB, and then fixed in 1% osmium tetroxide (Santa Cruz Biotechnology, Dallas, TX) in SCB for 1 hour at room temperature, which was followed by en bloc staining with 2% uranyl acetate, dehydration through graded alcohol and 100% acetone and embedment in Eponate (Ted Pella, Redding, CA). After at least 24 hours of polymerization at 56°C, 60 nm thick sections were cut with a diamond knife, counterstained with 5% uranyl acetate in methanol and in Reynold’s lead citrate for 10 min each, and then viewed with Jeol 100CX transmission electron microscope at 80 k. Digital images were collected with an XR40 Digital Camera (Advanced Microscopy Techniques Corp., Danvers, MA, USA).

### 2.5 Western blot analyses

CTL- and CLN6-IPSCs were homogenized in RIPA buffer (Sigma-Aldrich, St. Louis, MO) supplemented with 1x Protease inhibitor cocktail (ThermoFisher Scientific, Waltham, MA). 30ug total protein was denatured under reducing conditions in sample loading buffer (Bio-Rad, Hercules, CA) by boiling for 10 min at 98°C before loading onto a 4-20% gel (Bio-Rad, Hercules, CA), then transferred to PVDF 0.22 m membrane (Bio-Rad, Hercules, CA). The membrane was blocked for 1 hour with 5% bovine serum albumin (BSA; Sigma-Aldrich, St. Louis, MO) in TBS (Bio-Rad, Hercules, CA) supplemented with 0.5% Tween (Bio-Rad, Hercules, CA), followed by overnight incubation with primary antibodies at 4°C. The membrane was washed 3 times with TBS-T buffer and incubated at room temperature with secondary antibodies for 2 hours. After washing 3 times with TBS-T, the proteins were visualized using ECL Western blotting detection reagents (ThermoFisher Scientific, Waltham, MA). Primary antibodies: CLN6 (kind donation from Stellar lab), GAPDH (Santa Cruz Biotechnology, Dallas, TX). Secondary antibodies: HRP conjugated anti-mouse and anti-rabbit (GE Healthcare, Chicago, IL).

### 2.6 RNA expression and Real-time (RT)-qPCR

Total cellular RNA at Days N1 and N14 was isolated using Qiagen RNeasy Mini kit with DNase treatment (Qiagen, Germantown, md) to determine levels of TPP1 and PPT1 mRNA. Total RNA (1 μg) was used for cDNA synthesis using the Quantitate Reverse Transcription kit for cDNA Synthesis for PCR (Qiagen, Germantown, md). Real-time PCR was performed using SYBR Green Supermix (Bio-Rad, Hercules, CA).

Primers: TPP1-F:TCAGCAACAGAGTGCCCATT,TPP1-R:GCAGCCACGGGTTACATCAA, PPT1-F: CTGTAGATTCGGAGGACCGC, PPT1-R:TGGTTTGGAAGAGTTAGGGGC

The expression levels of respective genes were normalized to corresponding GAPDH values and are shown as fold change relative to the value of the control sample (ΔΔCt method). All sample analyses were carried out in triplicate.

### 2.7 Enzyme activity

TPP1 activity: 30mg of protein from scraped cells at Days N1 and N14 were incubated with 150ml of Acetate buffer (pH:4) supplemented with 10mM of protease inhibitors pepstatin A and E64 containing 62.5mM of AAF-MCA (Ala-Ala-Phe7-amido-4-methylcoumarin) (Sigma-Aldrich, St. Louis, MO). Fluorescence was measured using a Spectra Max spectrophotometer (Molecular Devices, San Jose, CA) at an extinction wavelength of 355nm and emission at 460nm.

Cathepsin D activity: 30mg of protein from scraped cells at day N1 and N14 were analyzed with the Cathepsin D Activity assay kit (ab65302; Abcam, Waltham, MA) following the manufacturer’s instructions.

### 2.8 Transcriptomics

Triplicate wells for each line were differentiated as above and split into cell pellets for either mRNA sequencing or protein analysis. Total RNA samples were assessed for concentration using a Qubit fluorometer (ThermoFisher Scientific, Waltham, MA) and quality using the 2100 Bioanalyzer (Agilent Technologies, Santa Clara, CA). Up to one µg of total RNA was purified for mRNA using the NEBNext^®^LPoly(A) mRNA Magnetic Isolation Module (New England Biolab Inc, Ipswich, MA). Stranded RNA-Seq library construction was performed using the IDT xGen Broad-Range RNA Library Prep Kit (Integrated DNA Technologies, Coralville, IA). Library concentration was measured with a Qubit fluorometer and library size on an Agilent 4200 TapeStation (Agilent Technologies). Libraries were multiplexed and sequenced on a NovaSeq 6000 (Illumina, San Diego, CA) using 75bp single-end sequencing. On average, approximately 30 million reads were generated from each sample.

Raw sequencing data were demultiplexed and converted to fastq format using bcl2fastq v2.20 (Illumina, San Diego, California). Then reads were aligned to the transcriptome using STAR (version 2.6.1) / RSEM (version 1.2.28)(Dobin et al., 2013; Li et al., 2014) with default parameters, using a custom human GRCh38 transcriptome reference downloaded from http://www.gencodegenes.org, containing all protein-coding and long non-coding RNA genes based on human GENCODE version 33 annotation. Expression counts for each gene in all samples were normalized by a modified trimmed mean of the M-values normalization method. Each gene was fitted into a negative binomial generalized linear model, and the Wald test was applied to assess the differential expressions between two sample groups by DESeq2 (version 1.26.0)(Love et al., 2014). Benjamini and Hochberg’s procedure was used to adjust for multiple hypothesis testing, and differential expression gene (DEG) candidates were selected with a false discovery rate of less than 0.05. DEG candidates were used for GO/KEGG enrichment analysis performed with ClusterProfileR (Wu et al., 2021).

## 3. Results

### 3.1 CLN6 Subjects had increased white matter signal abnormalities

Nine subjects were seen in either the NIH Intramural Undiagnosed Diseases Program (NIH-UDP)(Gahl et al., 2012; Gahl and Tifft, 2011) or the Cedars-Sinai Pediatric Neurogenetics Clinic (PNC). Each subject underwent record review and clinical evaluation. Some subjects had been reported previously (Chin et al., 2019). We evaluated each subjects’ previous neuroimaging to and found WMSAs were present on each subjects’ brain MRI, whether those neuroimaging studies were performed early or late in their disease course (Fig 1). WMSAs were predominantly present in the periventricular and occipital regions and were similar to previous case reports (Biswas et al., 2020; Cannelli et al., 2009; Guerreiro et al., 2013; Jadav et al., 2014; Katz et al., 2020; Rus et al., 2022; Sawiak et al., 2015; Sun et al., 2018; White et al., 2018). Brief summaries of each subject’s genotype, phenotype and white matter findings are presented in the Supplementary Information.

**Figure 1.**
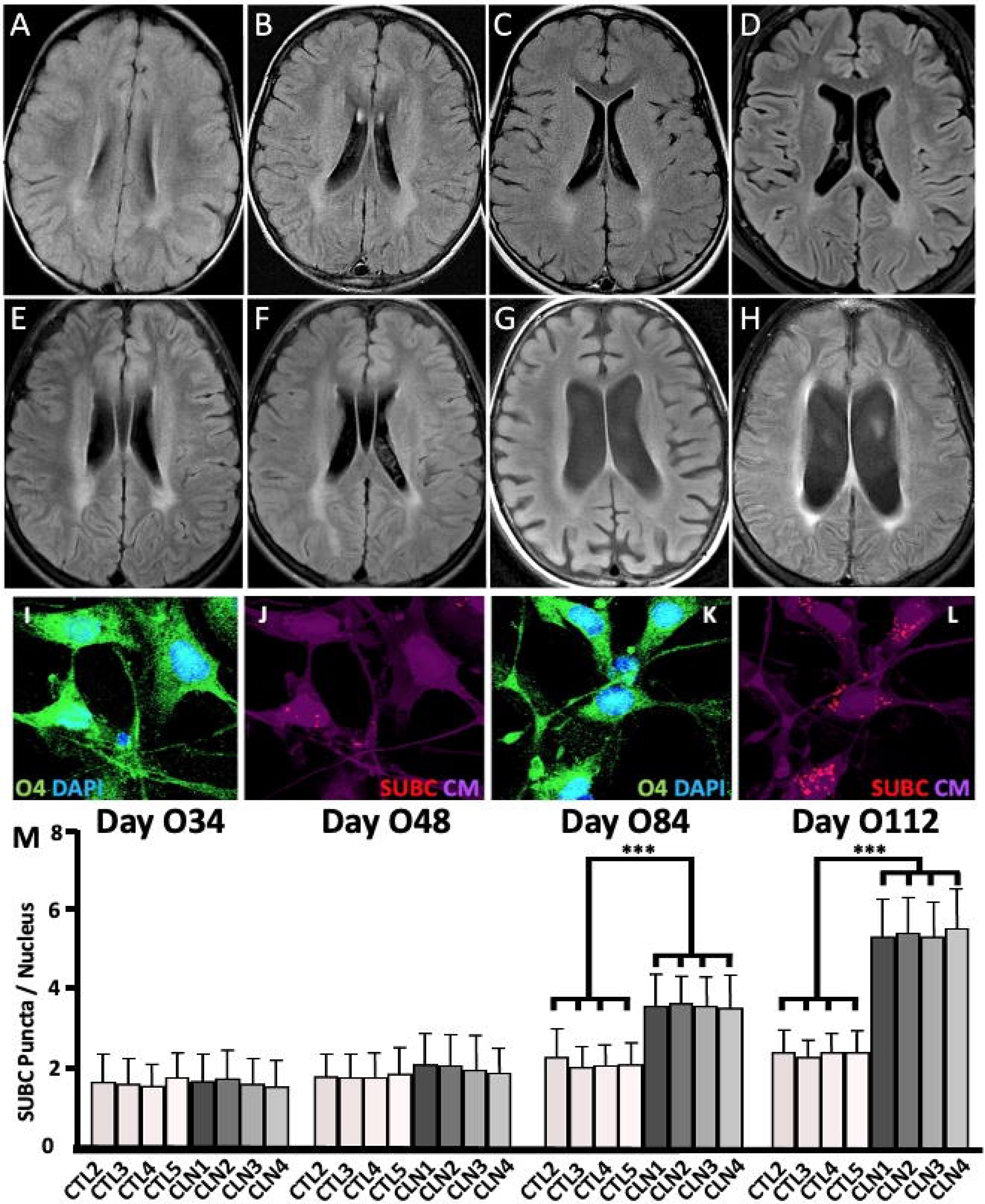
White matter and oligodendroglial changes in CLN6 cells. (**A-H**) Brain MRIs from multiple subjects reveal the presence of confluent bilateral supratentorial WMSAs involving the centrum semiovale and corona radiata often with extension into occipital white matter tracts (yellow arrows). (**A**) Subject 6 at 2 years, (**B**) Subject 5 at 5 years, (**C**) Subject 7 at 5 years, (**D**) Subject 4 at 11 years, (**E**) Subject 2 at 8 years (**F)** Subject 2 at 8 years (**G**) Subject 3 at ∼12 years (**H**) Subject 1 at 8 years. (**I-L**) Representative images show O4 marker expression/SUBC in Day O112 CTL-(**I/J**) and CLN6-(**K/L**) IPSC-derived Oligodendroglia. (**M**) Quantification of SUBC+ puncta in CTL (CTL2-5) and CLN6 (CLN1-4) IPSC-derived cells during oligodendroglial differentiation over time including Days O34, O48, O84 and O112 of differentiation. Values represent mean ± S.D.; N = 3 biological replicates per cell line; One-Way ANOVA Bonferroni’s multiple comparison test: *p < 0.05; ** p < 0.01; *** p < 0.001.

### 3.2 CTL and CLN6-IPSCs had similar characteristics

IPSCs were generated from peripheral blood mononuclear cells (PBMCs) from four CLN6 subjects (Subject 8 (CLN1), Subject 6 (CLN2), Subject 5 (CLN3), Subject 9 (CLN4)) along with five CTL subjects (Subjects CTL1-CTL5) (see Supplementary Information and Supplemental Table 1). CLN6-IPSCs did not express any recognizable CLN6 protein (Supplemental Fig. 1). CLN6- and CTL-IPSCs had no overt differences in ICC evaluation of several organelle markers (protein disulfide-isomerase (PDI), lysosomal associated membrane protein1 (LAMP1), Microtubule-associated protein 1A/1B-light chain 3 (LC3). CLN6- and CTL-IPSCs also had no overt differences or accumulation of SUBC^+^ ICC or Nile Red staining.

### 3.3 Increased SUBC^+^ storage CLN6 Oligodendroglia cultures

CLN6- and CTL-IPSCs were differentiated into oligodendroglia using well-established protocols to evaluate for storage accumulation in CLN6 patient-derived cells (Hu et al., 2009a; Hu et al., 2009b; Wang et al., 2013). Cells were cultured as neurospheres in suspension, were differentiated through several stages, and collected at representative time-points as follows: i) Pre-OPC stage (timepoint: Day O34) ii) OPC stage (timepoint: Day O48), iii) Immature Oligodendrocyte stage (timepoint: Day O84) and iv) Oligodendrocyte stage (timepoint: Day O112). ICC testing was performed for several oligodendroglial cell-type specific markers that included: Days O34 and O48: OLIG2 (pre-OPC marker) and NKX2.2 (OPC marker); Days O34, O48, O84, and O112: NG2 and PDGFRa (immature oligodendrocyte markers); and Days O84 and O112: O4 (oligodendrocyte marker). CTL- and CLN6-cells expressed similar levels of each marker throughout the oligodendroglial differentiation process with no significant expression level differences between the two genotypes (Supplemental Figs. 2-6). As expected, the OLIG2 signal was present in >80% of CTL- and CLN6-cells at both O32 and O48 time-points indicating that differentiation was moving forward appropriately. NKX2.2 signal was present in >50% at Day O34 and >80% of cells at Day O48, consistent with the maturation process. PDGFR-alpha signal was absent at Day O34 and present in very low amounts on Day O48 and, as expected, increased to involve >80% of cells on Days O84 and O112. Finally, NG2 and O4 were present in ∼80% of cells only on Days O84 and O112. SUBC^+^ storage material was significantly increased in the CLN6-as compared to CTL-lines on Days O84 and 0112 (Fig 1) p< 0.001 for both timepoints with one-way ANOVA Bonferroni’s multiple comparison test), but was not present in earlier stages indicating that storage accumulation was only present during the later terminal differentiation stages.

### 3.4 Increased SUBC^+^ storage in CLN6-neuronal cultures

CTL- and CLN6-IPSCs were also differentiated into excitatory cortical neurons using a dual-SMAD protocol (Shi et al., 2012b). IPSCs underwent neural epithelial induction and generated neural rosettes made up of IPSC-derived NPCs, which were expanded for several days until >80% confluency. As this could occur at different rates in different cell lines, we froze neural rosettes at this time and kept them at −80°C for short-term storage to better synchronize differentiation during the next steps of neuronal terminal differentiation (Supplemental Fig. 7). Once thawed, cultures were designated as “Day Post-Thaw 1” (Day PT1, subsequent days were designated Day PT2, PT3, etc.). NPCs continued to undergo neuronal terminal differentiation and were fixed and harvested at specific timepoints during this period (Days PT7, PT14, PT21, and PT28) for subsequent ICC and other staining procedures.

CTL- and CLN6-lines had no significant differences in Neuronal Nuclear Antigen (NeuN) expression. Less than 10% of lines were NeuN^+^ at Day PT7, which increased to greater than 80% of cells by Day PT14 and subsequent time points (Supplemental Fig. 8). CLN6-NPCs in neural rosettes did not have overt CLN6 pathology. CLN6-neuronal cultures did not have significantly increased SUBC^+^ storage on Day PT7, but were significantly increased at all time-points afterwards (Fig 2). Days PT14 and PT21 had the most significantly different amount of storage (0.5-0.8 versus 0.2-0.4 SUBC puncta per cell) (p < 0.001 for both time-points with one-way ANOVA Bonferroni’s multiple comparison test), while Day PT28 differences exhibited less significance (p < 0.05 and <0.01 with one-way ANOVA Bonferroni’s multiple comparison test).

**Figure 2.**
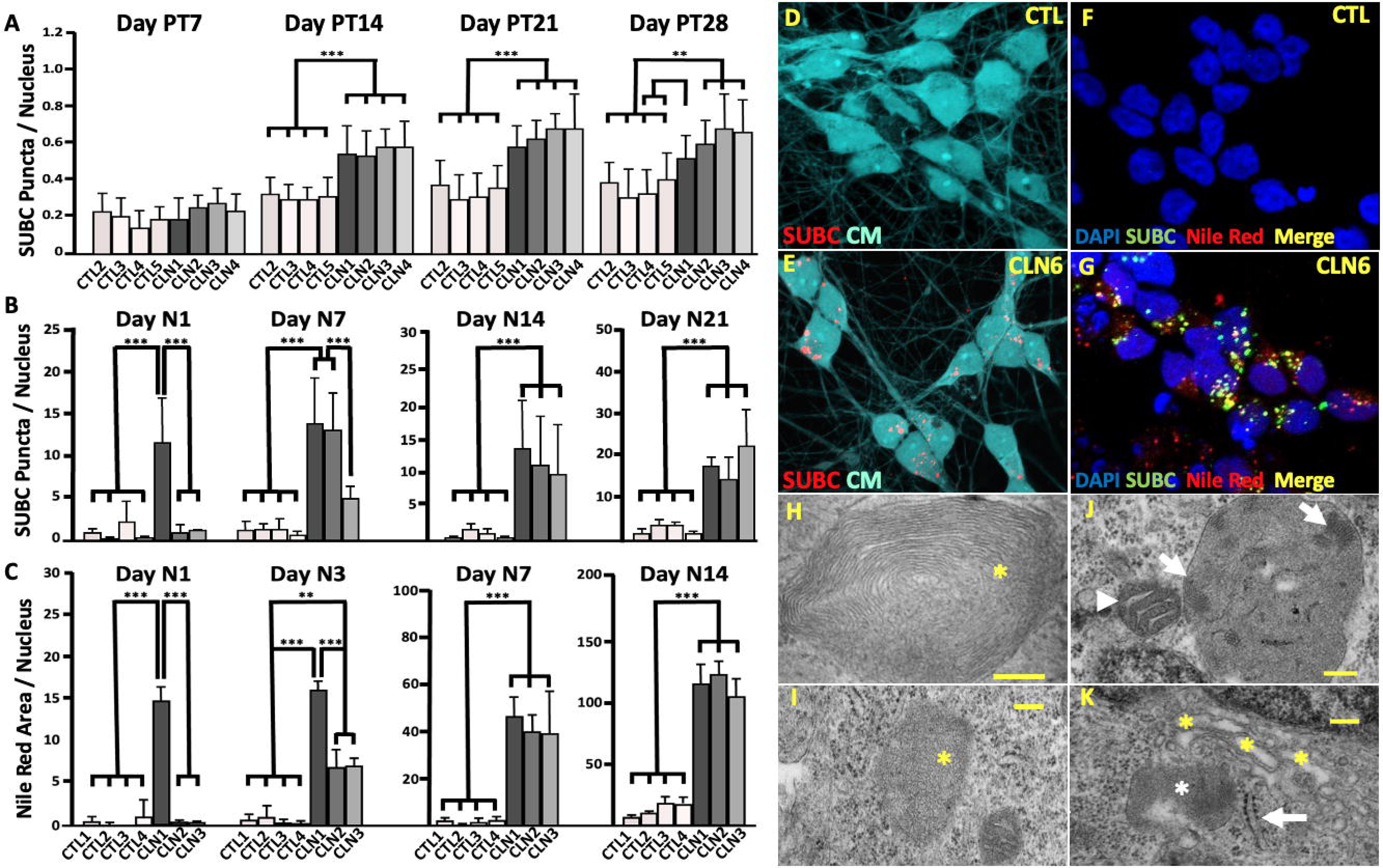
Storage accumulation in IPSC-derived neurons. (**A-C**) SUBC+ puncta and Nile Red+ area is increased over time in CLN6 (CLN1-4) versus CTL (CTL2-5) cell lines differentiated into neurons. (**A**) SUBC+ puncta accumulation in IPSC-derived cells during neuronal differentiation with dual-SMAD inhibition protocol, (**B**) SUBC+ puncta accumulation in I3N-neurons after treatment with DOX, (**C**) and Nile Red+ signal in I3N-neurons after treatment with DOX. Values represent mean ± S.D.; (**A**) N = 4 or (**B-C**) N = 3 biological replicates per cell line; One-Way ANOVA Bonferroni’s multiple comparison test: *p < 0.05; ** p < 0.01; *** p < 0.001 (**D-E**) Representative images showing the presence of SUBC+ puncta in (**D**) CTL- and (**E**) CLN6 cells (CM= cell mask stain). (**F-G**) Presence of SUBC+ puncta in (**F**) CTL- and (**G**) CLN6 cells (stained with DAPI, Nile Red, and SUBC ICC). (**H-K**) Ultrastructural analysis of CLN6-I3N-neurons (bars = 200nm) (**H**) CLN1 cell with fingerprint body (asterisk) (direct magnification 50,000X) (**I**) CLN2 cell with curvilinear body (asterisk)(direct magnification 25,000X) (**J**) CLN2 cell with inclusion with fragmented fingerprint features (arrows) and mitochondria with swollen cristae (arrowhead) (direct magnification 25,000X) (**K**) CLN2 cell with abnormal dilatation of Golgi (yellow asterisks), fingerprint body (asterisk), and slightly dilated rough ER (arrow)(direct magnification 25,000X).

### 3.5 Patient-derived I3N-IPSC characteristics

Drawbacks to using small molecule-based differentiation methods for evaluating differentiating and terminally differentiating cells include: i) the cultures containing heterogeneous cell populations ranging from undifferentiated to terminally-differentiated cells; ii) variable differentiation rates and ages of target cells (e.g., neurons); iii) batch-to-batch variability in the rate and amount of differentiation; iv) difficulty isolating targeted cells from the cultures; and v) the presence of mitotic and terminally-differentiated cells within the cultures (Zhang et al., 2013; Fernandopulle et al., 2018). Because of these issues, we utilized the I3N system for directed differentiation of I3N-IPSCs into I3N-neurons with a DOX-inducible NGN2 transgene. This technique uses clonal I3N-IPSC lines to generate clonal I3N-neuronal cultures, which produces synchronous terminally-differentiated cultures of I3N-neurons, allowing for more direct phenotypic evaluations without the confounding presence/influence of other undifferentiated neural cells (Fernandopulle et al., 2018; Wang et al., 2017).

Generation of I3N-IPSCs involved transfecting of parental IPSC lines with plasmids containing the inducible transgene and sequences for stable integration into the AAVS1 safe harbor locus using TALEN technology (Fernandopulle et al., 2018; Wang et al., 2017). The AAVS1 locus allows for continued expression of the transgene in IPSCs and during subsequent differentiation into neurons or other lineages. Clones were isolated, expanded, and then treated with DOX to screen for inducible neuronal differentiation by evaluating neuronal morphology and TUJ1^+^ immunostaining. Clones with ≥ 95% TUJ1^+^ cells in response to DOX were then tested for proper integration at the AAVS1 locus by PCR (Supplemental Fig 9)(Fernandopulle et al., 2018; Wang et al., 2017). In total, this time-intensive process generated four CTL-I3N-IPSC (from CTL1-4) and three CLN6-I3N-IPSC (from CLN1-3) lines that had proper integration into the AAVS1 site and normal karyotypes. When treated with DOX these I3N-IPSCs rapidly differentiated into TUJ1^+^ neurons.

### 3.6 Increased SUBC^+^ and Nile Red^+^ storage accumulation in CLN6-I3N-Neurons

All CTL- and CLN6-I3N-IPSC lines underwent DOX treatment for 3 days (Days D1-D3) and then were cultured for up to 21 days (Day N1-N21). Cultures were harvested on Days N1, N7, N14, and N21, evaluated with ICC for SUBC, and co-stained with Cell Mask and DAPI (Fig 2). By Day N1, >95% of cells in all I3N-lines exhibited neuronal morphology, and the TUJ1^+^ signal was consistent with an immature neuron identity. Significant increased SUBC^+^ storage was present on Day N1 in line CLN1, with both CLN1- and CLN2-I3N lines having significant SUBC^+^ storage by Day N7, with all three CLN6-I3N lines having significant SUBC^+^ storage by Day N14 that was still present at Day N21 (Fig 2). CLN-I3N-neuronal cultures had >10 SUBC^+^ puncta/nuclei compared to only ∼0.5 SUBC^+^ puncta/nuclei seen in CLN6-IPSC cultures differentiated with the dual-SMAD inhibition protocol.

Nile Red staining is an alternative method of identifying and quantifying cellular storage (Greenspan et al., 1985; Sima et al., 2018). Nile Red dye accumulates in cellular lipids/proteolipids after short incubation and subsequent wash steps. This method is cheaper and faster than SUBC^+^ ICC and may be an attractive storage detection method for use in upscaled automated assays. We evaluated CTL- and CLN-I3N cultures with Nile Red staining on Days N1, N3, N7, and N14 to determine if this method would be valuable for storage quantification in this context. Interestingly, Nile Red+ storage was present earlier than SUBC^+^ storage with CLN1-I3N-neurons having significant Nile Red^+^ storage by Day N1 and all three CLN-I3N lines having significant Nile Red^+^ storage by Day N3 (Fig 2). In order to determine whether Nile Red^+^ and SUBC^+^ signals co-localize in I3N-neurons, Day N3 CTL- and CLN-I3N-cultures were stained with both methods. Little to no Nile Red^+^ or SUBC^+^ signal was present in CTL-I3N-cultures. CLN-I3N-cultures had abundant Nile Red^+^ and SUBC^+^ signals, with Nile Red^+^ signals being more abundant than SUBC^+^ signals. SUBC^+^ was always seen in close proximity to the Nile Red^+^ signal (Fig. 2).

### 3.7 Abnormal ultrastructural features in CLN6-I3N-Neurons

Several ultrastructural findings are common in CLN6 and so I3N-neurons were evaluated by transmission electron microscopy to determine whether these ultrastructural changes were also seen in I3N-neurons. CTL- and CLN-I3N-neurons (CTL2, CLN1 and CLN2, respectively) were collected on Day N7 and imaged with transmission electron microscopy. Both CLN-I3N-neurons had numerous examples of ultrastructural pathology previously seen in CLN6 that included cytoplasmic inclusions with mixed fingerprint and curvilinear bodies, cystic Golgi enlargement, and abnormal megaconial mitochondria with loss of cristae (Fig 2).

### 3.8 Golgi enlargement in CLN6-I3N-Neurons

The morphology and size of several organelles were also evaluated to identify differences between the CTL- and CLN6-I3N-neurons. I3N-neuron samples were isolated on Days N1-N14 and evaluated with ICC for markers of the endoplasmic reticulum (PDI/BiP), mitochondria (GRP75) and cis-Golgi (GM130) and SUBC. Preliminary studies revealed no change in PDI/BiP or GRP75 immunoreactivity (data not shown), although GM130 significantly differed between the CTL- and CLN6-I3N-neurons at Day N14 (Fig 3). CTL-I3N-cells’ Golgi were juxtanuclear in position and morphologically compact, while the CLN6-I3N-cells’ Golgi had a more expanded morphology associated with a larger Golgi/soma area ratio (Fig 3). This trend was somewhat present on Day N1, but was significant on Day N14. GM130 and SUBC immunosignal did not co-localize in these cells (Fig 3).

**Figure 3.**
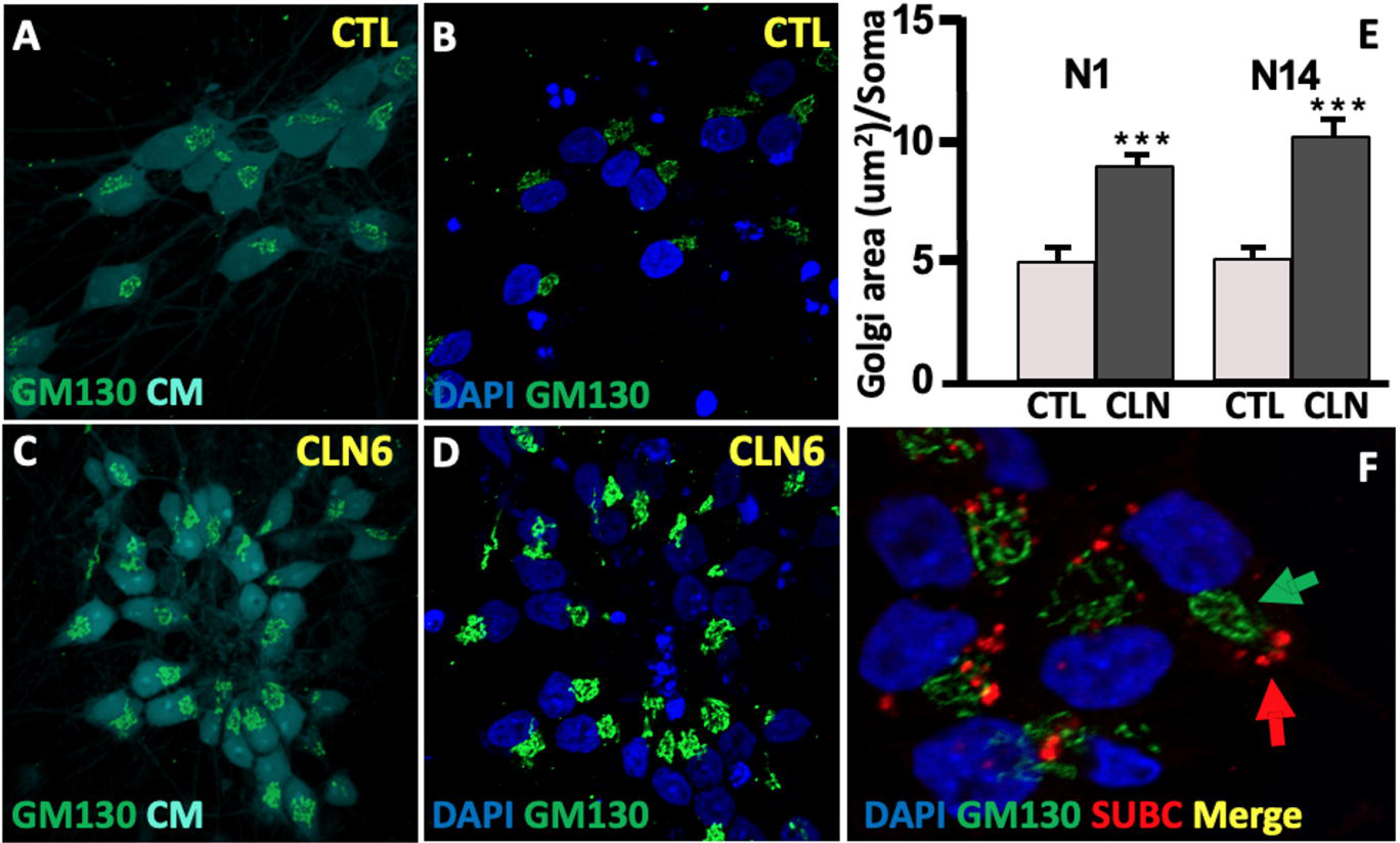
Golgi enlargement in CLN6 cells. (**A-D**) Representative images for (**A-B**) CTL and (**C-D**) CLN6 neurons stained for (**A/C**) GM130 and Cell Mask (CM) or (**B/D**) GM130 and DAPI. (**E**) Golgi body area was significantly increased in CLN6-I3N-neurons at Day N14. Data points presented as means of ≥4 replicates from each individual cell line, with bars presenting overall group means and standard deviations, *p < 0.05 from mixed model regression. (**F**) Representative images for CLN6-I3N-neurons stained against GM130 and SUBC indicate that the two markers do not co-localize.

### 3.9 Increased LAMP1^+^ size and altered localization in CLN6 I3N-Neurons

Lysosomal size and distribution were the next parameters to be evaluated. CLN6-I3N-neurons had significantly increased LAMP1 signal at Day N14 (Fig 4). As expected, SUBC^+^ signal was almost always present within the LAMP1 signal (Dunn et al., 2011). In addition, CLN6-LAMP1^+^ puncta were larger and dispersed throughout the cell bodies with large amounts of signal extending into neuritic processes, while LAMP1^+^ puncta in CTL-cells appeared smaller and primarily located in the perinuclear region of the cell soma (Fig 4). These findings were consistent with other studies involving other NCL-associated membrane-bound proteins and may indicate the presence of axonal and synaptic dysfunction in those NCL disorders (Lojewski et al., 2014).

**Figure 4.**
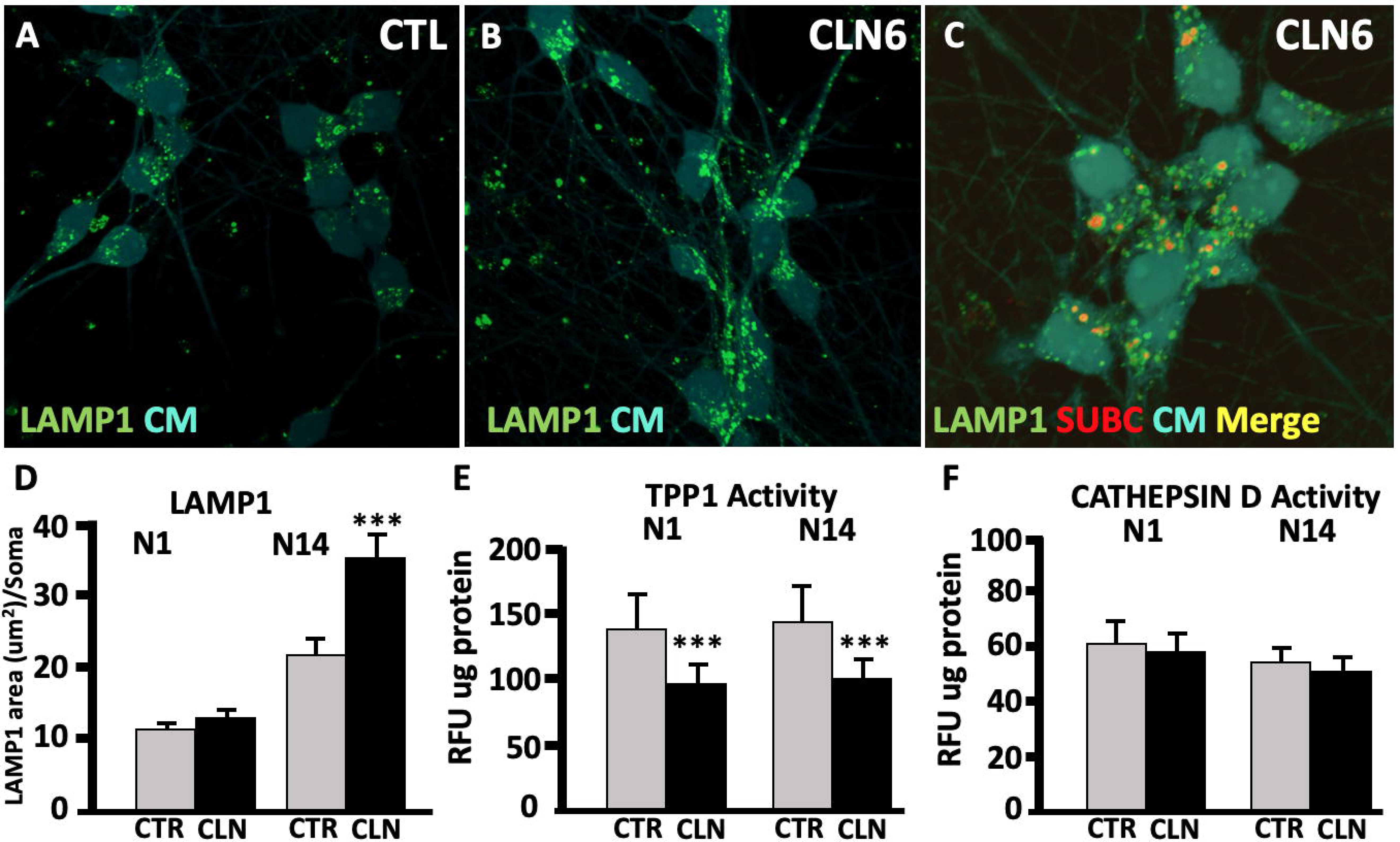
CLN6-I3N-neurons have increased LAMP1+ signal and decreased TPP1 activity. (**A-B**) Representative images for (**A**) CTL- and (**B**) CLN6-I3N-neurons stained for LAMP1 and Cell Mask. (**C**) Representative images for CLN6-I3N-neurons stained for LAMP1, SUBC, and Cell Mask reveal SUBC^+^ puncta co-localize with LAMP1^+^ signal. (D) LAMP1+ area is increased in CLN6-I3N-neurons on Day N14, but not on Day N1. Data points presented as means of ≥ 6 replicates from each individual cell line, with bars presenting overall group means and standard deviations, *p < 0.05 from mixed model regression. (E) TPP1 enzymatic activity is decreased in CLN6-I3N-neurons, data points presented as means of 2 replicates from each individual cell line, with bars presenting overall group means and standard deviations, *p < 0.05 from mixed model regression. Data log-transformed prior to analysis. (F) Cathepsin D activity was unaffected. Data points presented as means of 2 replicates from each individual cell line, with bars presenting overall group means and standard deviations, *p < 0.05 from mixed model regression. Data log-transformed prior to analysis. Analysis also corrected for significant variation across date of assay.

### 3.10 Abnormal lysosomal acid hydrolases Activity in CLN6-I3N-Neurons

Functional enzyme assays of tripeptidyl peptidase-1 (TPP1) and Cathepsin D (CTSD) activity were performed on I3N-neurons harvested at Days N1 and N14. CTSD activity was not significantly different between CTL- and CLN6-I3N-cells at either timepoint; however, TPP1 activity was significantly decreased in CLN6-I3N-Neurons on both Day N1 and N14 (Fig 4). Transcriptional evaluations of TPP1, CTSD, and palmitoyl-protein thioesterase-1 (PPT1) with quantitative reverse-transcriptase-mediated polymerase chain reaction (qRT-PCR) at Days N1 and Day N14 revealed no significant differences between CTL- and CLN6-I3N-neurons, indicating that decreased TPP1 enzymatic function was not due to downregulation of *TPP1* gene expression (Supplemental Fig 10).

### 3.11 Transcriptomic differences between CTL- and CLN6-I3N-Neurons

Bulk transcriptomic analyses of I3N-Neurons were performed to evaluate differential gene expression between three CLN6- and four CTL-I3N-neuron lines. Cells were differentiated and harvested at Day N5 to provide an early window on cellular changes after the accumulation of significant Nile Red^+^ storage in all lines at Day N3. RNA was collected for transcriptomic evaluation to identify differentially expressed genes (DEGs) in CLN6-I3N-neurons (adjusted p-value < 0.05 (aka Benjamini Hochberg corrected)). PCA maps of transcriptomic data had appropriate clustering of within CLN6-I3N-neuron and CTL-I3N-neuron lines, but were divergent between CLN6 and CTLs (Supplemental Fig 11). Comparative expression analysis identified 1382 total DEGs in CLN6 cells, with 645 being significantly upregulated and 737 significantly downregulated (Fig 5, Supplemental Fig 11). Gene Ontology (GO) and Canonical Pathway analyses identified numerous significant pathways and biological processes (adjusted p-value < 0.05) (Fig 5D). Many of those processes are linked to organelle localization, microtubule transport, axonal maintenance, synaptic formation, and neuronal apoptosis.

**Figure 5.**
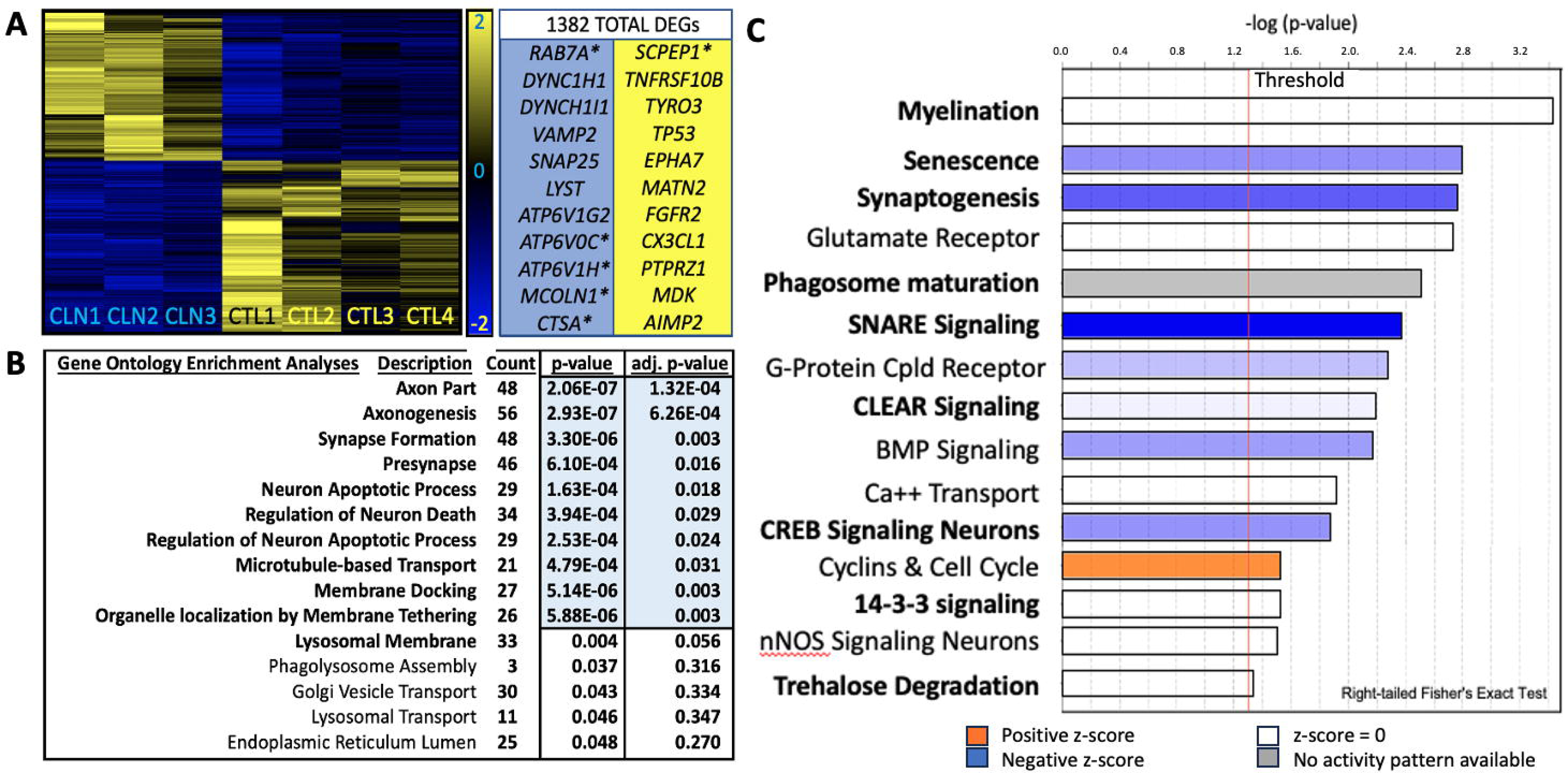
Bulk RNA-Seq in I3N-Neurons at Day N5. (**A**) Representative Heat Map with list of some representative DEGs (blue column shows down-regulated DEGs, yellow column shows up-regulated DEGs (*CLEAR Network genes). (**B**) Gene Ontology (GO) Enrichment Analysis showed several GO groupings with significant differences (adj p value < 0.05) and other GO groupings of interest **C**) Canonical Signaling Pathway Analysis had significantly affected groupings that included terms involved in Synaptogenesis, SNARE Signaling, Phagosome maturation & CLEAR Network Signaling.

Several of the 1300+ DEGs (see Supplemental Table 2 for adjusted p-values) identified encoded proteins involved in lysosomal function, size, numbers, and reformation (*LYST, MCOLN1*, *SCPEP1, CTSA, SMPD1*). For example, LYST functions in trafficking lysosomes and regulating lysosomal fission/fusion (Serra-Vinardell et al., 2023), MCOLN1 encodes mucolipin-1, a cation channel associated with Mucolipidosis IV (Sun et al., 2000), while CTSA and SMPD1 are acid hydrolases involved in galactosialidosis and Niemann-Pick types A/B respectively (Levran et al., 1991; Pan et al., 2014). Other lysosome-specific DEGs included several that encode lysosomal-vesicular proton pump subunits (*ATP6V1G2*, *ATP6V1H, ATP6V0C*)(Li et al., 2020; Mattison et al., 2023). Non-lysosomal DEGs encoded genes involved in cellular motor subunits (*KIF3C, DYNC1H1*, *DYNC1I1*)(Hirokawa et al., 2010), trafficking proteins (*RAB7A, VPS39, STXBP1*)(Caplan et al., 2001; Hamdan et al., 2011; Meggouh et al., 2006), or are involved in synaptic function (*VAMP2, SNAP25, SLC17A7, ID4*)(Rechtzigel et al., 2022). Additional identified DEGs encode cytokines and cytokine receptors (*CX3CL1, FGFR2, EPHA7, MDK*)(Chen et al., 2017; Depaepe et al., 2005; Nakamura et al., 1998; Subbarayan et al., 2022), along with several genes involved with neuronal apoptosis (*C5AR1, PINK1, TYRO3*)(Brunelli et al., 2022; Mahajan et al., 2016; Zhu et al., 2016).

GO Enrichment analysis revealed numerous GO terms that were significantly different in gene expression in CLN6-compared to CTL-cells (Fig 5). These GO terms included several involved in axon and synapse formation and maintenance (Axon-Part (GO:0033267); Axonogenesis (GO:0007409); Negative-Regulation-of-Neuron-Projection-Development (GO:0010977); Synapse-Organization (GO:0050808); and Neuron-to-Neuron-Synapse (GO:0098984)), as well as several GO terms involved in neuronal apoptosis (Neuron-Apoptotic-Process (GO:0051402); Regulation-of-Neuron-Death (GO:1901214). Other relevant and significantly altered GO terms included terms involved in intracellular trafficking of endo-lysosomal components (Endocytic-Vesicle (GO:0030139), Organelle-Localization-by-Membrane-Tethering (GO:0140056); and Membrane-Docking (GO:0022406)). Several other GO terms that included specific lysosomal and Golgi-centric groups were also altered, but had an adjusted p-value > 0.05 (Lysosomal-Membrane (GO:0005765; p<0.003, adj. p-value <0.082, fold enrichment count: 1.641), Golgi-Vesicle-Transport (GO:0048193; p<0.043, adj. p-value <0.334, fold enrichment count: 1.391) and Lysosomal-Transport (GO:0007041; p<0.046, adj. p-value <3.47, fold enrichment count: 1.424)). Canonical Pathway Analysis indicated that multiple pathways were significantly different in CLN6-I3N-neurons that included: Phagosome-Maturation, Glutamate-Receptor, Myelination, SNARE-Signaling, and Synaptogenesis-Signaling (Fig 5). Interestingly, Canonical Pathway analysis also indicated the CLEAR signaling pathway was significantly down-regulated at Day N5 in CLN6-I3N-Neurons indicating this pathway may have entered the chronic stimulation phase (Fig. 5)(Palmieri et al., 2011).

## 4. Discussion

Our goal for this study was to discover and report new insights into the clinical, cellular, and molecular pathology of CLN6. Clinically, we discussed the prevalence of WMSAs in our genetically-confirmed CLN6 patient cohort and the potential role oligodendroglia may play in CLN6. We also generated renewable patient-derived IPSC-models of CLN6 to model disease pathology in different neural cell types, including oligodendroglia and neurons. CLN6- and CTL-IPSCs were further engineered to contain the DOX-inducible I3N-system for rapid directed differentiation of I3N-IPSCs to I3N-neurons. Our initial studies with IPSCs used SUBC^+^ as a biomarker for disease and showed SUBC+ storage accumulation occurred in both IPSC-derived CLN6-oligodendroglia and-neurons. Subsequent work in I3N-neurons also used Nile Red^+^ staining as an additional biomarker to detect lipid/proteolipid storage accumulation. Electron microscopy confirmed the presence of ultrastructural changes commonly seen in the CLN6 were also present in IPSC-derived CLN6-I3N-neurons. Subsequent analysis of CLN6-I3N-neurons revealed the presence of changes in the structure and size of lysosomes and Golgi organelles, as well as decreased TPP1 enzymatic activity. Further bulk-transcriptomic analyses identified variations in the transcriptional expression of numerous gene pathways involved in axonal, synaptic, and neuronal apoptotic processes. Interestingly, many of these transcriptional changes were correlated with shifts in transcript and protein expression that had been identified in other models of NCL and lysosomal disease (Gomez-Giro et al., 2019; Pérez-Brangulí et al., 2014; Rechtzigel et al., 2022; Sun et al., 2000). These data indicate that our CLN6-IPSCs and their derivatives can serve as appropriate human cellular models for the CLN6. Of course, there are limitations to using IPSCs as models for disease, but overall, our initial data indicate that we have generated a novel renewable cellular model of CLN6 that can assist with initial in vitro investigations for better understanding CLN6’s pathology and its progression, as well as being a useful tool in initial screenings of potential therapeutic interventions for this difficult disease.

CLN6A is a childhood disorder that is known to cause progressive dementia, motor decline, visual loss, and seizures. For the most part, the NCLs are thought to be disorders of grey matter, but several case reports have noted some CLN6 patients presented with white matter changes. Here, we report a nine-subject cohort from our undiagnosed diseases clinics with CLN6A clinical phenotypes. This cohort had three sibling pairs with five females and four males. There were eight variants: seven of which had been previously reported, along with one novel variant (c.655 +1 G>A; IVS6+1) (Andrade et al., 2012; Badilla-Porras et al., 2022; Berkovic et al., 2019; Chin et al., 2019; White et al., 2018). Four subjects possessed homozygous variants: with two of those being siblings. Interestingly, neuroimaging of this cohort showed the typical grey matter findings associated with CLN6, but all subjects were found to also express WMSAs (Fig 1). WMSAs have been previously reported in several CLN6 case reports, but have not commonly been considered part of the disorder. Recently, WMSAs have been reported in several animal models of CLN6, which indicate white matter involvement may be common in CLN6 compared to other NCLs (Biswas et al., 2020; Cannelli et al., 2009; Faller et al., 2015; Johnson et al., 2020; Katz et al., 2020; Sun et al., 2018; Thelen et al., 2012a; Thelen et al., 2012b). This may be why CLN6 has been the most common NCL referred to our undiagnosed patient clinics, as the presence of WMSAs/leukodystrophy may confound the interpretation of genetic testing due to this not being understood as part of the CLN6 presentation. These results indicate that future clinical studies of CLN6 should evaluate whether WMSAs are a consistent or variable part of the human CLN6 phenotype.

Due to our clinical neuroimaging data, we wanted to determine whether oligodendroglia could accumulate storage material in CLN6. CLN6- and CTL-IPSCs were differentiated using established oligodendroglial protocols for over 112 days. Cells were routinely tested for SUBC accumulation and the chronological expression of differentiation markers that included OLIG1, NKX2.2, PDGFRa, NG2, and O4. No differences were seen in expression levels of these differentiation markers in CTL- and CLN6-cells, indicating that CLN6 did not inhibit oligodendroglial differentiation. At Days O84 and O112, the O4 marker was present in >80% of all cell lines, indicating that a majority of cells were terminally differentiated IPSC-derived oligodendroglia cells. SUBC accumulation was low in CTL- and CLN6 cells during the early stages of the protocol, but CLN6-cells had a significant increase in cytoplasmic SUBC^+^ signal at Days O84 and O112. These data indicate that SUBC^+^ signal did not accumulate during the early mitotic stages of the differentiation process, but was present during stages of terminal differentiation. The data also indicated that oligodendroglia could accumulate SUBC^+^ storage and may be involved in the generation of WMSAs. The data also suggest that any therapeutic interventions for CLN6 may also require the additional targeting of white matter for efficacy (Cain et al., 2019; Geraets et al., 2016).

Our CTL- and CLN6-IPSCs were also differentiated using well-established neuronal protocols (dual SMAD inhibition; Shi et al., 2012a; Shi et al., 2012b) that were modified to allow for better synchronization of lines by freezing NPC cultures early (∼15-20 days) when >80% of the cells had reached the neural rosette stage. When appropriate, cultures were then thawed to produce neurons over the next four weeks (Supplemental Fig. 8). CTL- and CLN6-cultures had similar percentages of NeuN^+^ cells present at every timepoint, indicating that neuronal differentiation was occurring at similar rates. At Day PT14, SUBC^+^ storage material was significantly increased in CLN6-cultures, which continued through Days PT21-PT28. The average number of SUBC^+^ puncta per cell was less than one, which indicated that not every cell contained SUBC^+^ storage. This was most likely due to the continued presence of mitotic cells in the cultures that had not yet undergone terminal differentiation (e.g., IPSCs, NPCs) that could dilute the level of accumulating storage with each continued cell division. Overall, our data indicated that IPSC-derived CLN6-neurons accumulated SUBC^+^ storage similar to their *in vivo* counterparts.

The use of the dual-SMAD inhibition protocols for neuronal differentiation had several limitations, including: i) a relatively prolonged period for generating neurons; ii) the heterogeneous nature of the neuronal cultures; iii) the heterogeneous age of the neurons in each culture; and iv) the lack of storage material in every cell. Because of these limitations, we utilized the I3N-system to generate a more flexible and precise neuronal models. The I3N-system uses a transgene that is targeted to the AAVS1 safe-harbor site. This transgene possesses a DOX-inducible NGN2 expression system that when activated by DOX-treatment rapidly differentiates I3N-IPSCs into I3N-neurons (Fernandopulle et al., 2018; Wang et al., 2017). We generated three CLN6- and four CTL-I3N-IPSC clonal lines that had proper AAVS1 site integration and normal karyotypes. DOX-treatment caused the majority (>95%) of these I3N-IPSCs to rapidly differentiate into TUJ1^+^ I3N-neurons within 1-3 days. One line had significantly increased SUBC^+^ puncta on Day N1 (CLN1 line), while all three lines had significantly increased SUBC^+^ storage by Day N14. In contrast to IPSC-derived neuronal cultures differentiated with dual SMAD inhibition protocols, each I3N-neuron possessed multiple SUBC^+^ puncta on average (5-15). Electron microscopic evaluation of CLN6-I3N-neurons revealed the presence of ultrastructural changes consistent with in vivo CLN6 findings that included rectilinear, fingerprint and curvilinear profiles associated with multi-lamellar structures. Subsequent evaluations with Nile Red staining revealed that lipid/proteolipid storage accumulated even faster than ICC-detected SUBC^+^ storage material, with all I3N-CLN6 lines having significantly increased Nile Red^+^ signal by Day N3. When CLN6-I3N-neurons were co-stained with Nile Red and SUBC-ICC, Nile Red^+^ signal was seen more prevalent than the SUBC^+^ signal, with most SUBC^+^ signals seemingly co-localizing with Nile Red^+^ signals. These results suggest that Nile Red staining may provide a simpler and earlier detection method for CLN6 storage compared with SUBC ICC, which may be particularly useful in automated therapeutic drug screening methods. Overall, these data indicate that I3N-neurons recapitulate storage features typically seen with *in vivo* CLN6 samples and can be used as a valid and renewable source of human cellular CLN6 modeling.

In addition to accumulating storage, CLN6-I3N-neurons had significantly altered morphological and enzymatic function. For example, significantly increased Golgi size (GM130^+^) was present at Day N14, although it did not co-localize with SUBC^+^ storage. Further study will be required to learn the specific causes of this phenomena, but this could be the result of organellar stress and protein accumulation in the absence of proper CLN6 protein sorting in the ER (which was not enlarged). Other NCLs and lysosomal disorders have also been shown to have abnormal Golgi phenotypes (Lojewski et al., 2014), whether these phenomena are related will require further study. In addition to Golgi abnormalities, lysosomal (LAMP1^+^) area was also increased in CLN6-I3N-neurons at Day N14. LAMP1^+^ signal was mostly seen within the cell bodies of CTL-I3N-neurons, while CLN6-I3N-neurons had extensive amounts of LAMP1^+^ signal that also extended into their neuronal processes. As with other NCLs, this could indicate lysosomal mis-trafficking or downstream abnormalities in axonal transport (Lojewski et al., 2014). SUBC^+^ storage co-localized with LAMP1^+^ signal in CLN6-I3N-neurons. Further evaluation of lysosomal function indicated that there were no differences in transcript levels of TPP1, PTT1, or CTSD at Days N1 and N14; however, TPP1 enzymatic activity was significantly decreased in CLN6-I3N-neurons at Days N1 and N14, while CTSD was normal at all time-points. This result was slightly unexpected, as both TPP1 and CTSD play a role in SUBC catabolism and CLN6 function. Why CTSD activity was normal in CLN6-I3N-neurons, but abnormal in mouse liver cells will require further evaluation (Bajaj et al., 2020; Gao et al., 2002).

The next step in our studies was to use bulk-transcriptomic analyses to identify differential gene expression between CLN6- and CTL-I3N-neurons. To do this, I3N-neurons were harvested at an early time-point (Day N5) to determine transcriptomic changes during early storage accumulation. PCA maps had appropriate divergent clustering between CLN6-I3N-neurons and CTL-I3N-neurons, indicating that the two populations were behaving similarly within each genotype, but differently between CTL- and CLN6-cells. Transcriptomic data also indicated that >1350 total DEGs were identified between CLN6- and CTL-I3N-neurons, which included genes encoding proteins involved in lysosomal generation/function, as well as genes involved with cellular trafficking, axonal generation, synaptic function, and neuronal apoptosis. Gene Ontology (GO) Enrichment analysis revealed ∼145 GO terms that were significantly different between CTL- and CLN6-lines. These GO terms included groups of genes involved in endocytic and lysosomal function, organellar transport, axonal generation, synapse function, and neuronal apoptosis. Canonical Pathway Analysis also revealed multiple pathways that were significantly altered, including pathways involved in phagosome-maturation, myelination, glutamate-receptor, SNARE-signaling, synaptogenesis-signaling, and CLEAR-signaling (Fig. 5)(Sardiello et al., 2009; Serra-Vinardell et al., 2023).

It is important to note that many DEGs and pathways altered in our CLN6 lines were also affected in other research involving lysosomal disorders (e.g. CLN3, CLN6, CLN8, SPG11)(Gomez-Giro et al., 2019; Pérez-Brangulí et al., 2014; Rechtzigel et al., 2022; Sun et al., 2000). In one informative study, the cerebral cortices of *Cln3^Dex7/8^*, *Cln6^nclf^*, and *Cln8^mnd^*null mice abnormally expressed several proteins involved in presynaptic function and vesicular acidification/fusion (Atp6v1h, Munc18, Vamp2)(Rechtzigel et al., 2022), which were also transcriptionally down-regulated in our human CLN6-I3N-neurons (*ATP6V1H*, *STXBP1* (MUNC18), and *VAMP2)*. ATP6V1H was significantly altered in all three mice, while MUNC18 and VAMP2 protein levels were specific to the CLN6^nclf^ mouse, indicating CLN6-specific overlap in our transcriptional and their protein results. VAMP2’s partner, *SNAP25,* also had reduced transcription in CLN6-I3N-neurons. Other proteins that were significantly decreased in these three mouse models included STX1A, STX1B, and ATP6V0D; however, in CLN6-I3N-neurons these genes were downregulated at a level just short of significance (adj p-value = 0.085, 0.060, and 0.075, respectively). In a different study using IPSC-derived CLN3-cerebral organoids, several axonal and synaptic proteins with abnormal expression were identified as DEGs (*RAB7A*, *SLC17A7*, ID4) that were also dysregulated in our transcriptional data (Gomez-Giro et al., 2019). Expression changes in CLN6 were also shared in other lysosomal diseases like SPG11, a storage disorder with NCL features that include cognitive decline, motor dysfunction, and retinal abnormalities (Pérez-Brangulí et al., 2014). IPSC-derived SPG11 neurons also had *VAMP2* as a DEG, along with two anterograde motors, *KIF3A* and *KIF5A*. Separate transport motor genes were differentially expressed in CLN6-I3N-neurons, including *KIF3C*, an anterograde motor, along with two dynein genes, *DYNC1H1* and *DYNCH1I1* (adj p-value = 0.019, 5.742 E-06, and 0.002, respectively). These overlapping results indicate that the pathology in NCLs and other lysosomal disorders may have substantial similarities that are not limited to the endosomal-lysosomal pathway, but include axonal, synaptic, and other organellar systems. Additional research will be important in identifying which of these systems may be the key driver of neuronal dysfunction in these disorders and their associated neurodegeneration.

Our data indicate that CLN6-I3N-IPSCs are valid, renewable, and accurate models for investigating CLN6 and its pathology. We believe these cells represent a valuable addition to previous *in vitro* CLN6 neuronal models, which often include primary neural cells derived from CLN6 animal models for data generation. Although informative, these primary neural cell models had significant limitations that included difficulty with cell isolation, variability between preparations, genetic manipulation, and the potential inadequacy of recapitulating human disease. Alternatives to primary neural cells include primary non-neural cell models from CLN6 animal models (Bajaj et al., 2019), although how well these non-neuronal cells can model mechanisms of CNS disease is difficult to determine (Bajaj et al., 2019). Human cellular models of CLN6 have been limited to mitotic fibroblasts and lymphoblasts that do not express substantial pathology and may be limited in utility (Cao et al., 2011; Heine et al., 2004). Our hope is that CLN6 research can utilize the alternative advantages of our CLN6-IPSC models.

In summary, we have generated new human cellular models of CLN6 that revealed new insight into CLN6’s effects on neurons and glia. We have shown that CLN6-IPSCs can generate pathologic neurons and glia and that CLN6-I3N-neurons share similarities with these and other *in vitro* and *in vivo* models. Some of the next steps in our work will be serialized transcriptomic evaluations of I3N-neurons at earlier and later time-points to better understand the cascade of abnormal gene regulatory signals that occur during storage accumulation. We will also further evaluate many of the DEGs identified in this and future studies to determine which may play major roles in neuronal dysfunction. Our hope is that this future work can identify potential targets for small molecule or anti-sense oligonucleotide therapies that target these downstream effects. Other tactics can include targeting I3N-IPSCs or I3N-neurons with viral vectors to express CLN6p at different time-points before or after the development of pathology to determine temporal windows for preventing or rescuing diseased neurons. Finally, CLN6-I3N-neurons may be most valuable for future large-scale therapeutic drug screening studies to identify any potential small molecule interventions that can be used to treat this difficult disease.

## Acknowledgments

The authors thank all the families with *CLN6* variants for their participation and involvement in this work. We thank the subjects and their families for their help, for providing medical information, and for unconditional and special care for their children. We also would like to extend special thanks to the Gray Foundation for their assistance and continued cooperation. We also want to thank Stella Lee, PhD, for gift of the anti-human CLN6 antibody. We would also like to thank Jonathan Mink, Jill Weimer, Jonathan Cooper, and Stephanie Hughes for discussions and assistance. We also acknowledge Carolina Diaz, Erica Caro, and Carmela Brito for clinical and technical assistance. We acknowledge Alexander Laperle, Ritchie Ho, Nur Yucer and Veronica Garcia for their assistance with this work. We are grateful to Barrington Burnett for critical review of the manuscript. The content is solely the responsibility of the authors and does not necessarily represent the official views of the NIH or Cedars-Sinai Medical Center. We are especially grateful for the exceptional and continued support from the Cedars-Sinai Fashion Industries Guild.

## Author contributions

MGO, JK, YKK, FDN, CB, DRA, CT and TMP designed and conceptualized the project. MGO, JK, YKK, AR, FDN, CS, SIB, HO, HMUF, MB, CF, LD, SM-A, RM-L, DRA, CT, CJT, and TMP performed experiments and collected data. MGO, JK, YKK, FDN, HO, HMUF, and MB maintained and differentiated IPSCs. MGO, JK, YKK, AR, FDN, HO, CB, LD, SM-A, RM-L, DRA, CJT, CT, NK, WAG, NS and TMP analyzed and/or graphed/illustrated data. MGO, JK, YKK, CB, HMUF, CF, SM-A, DA, CT, CJT, NK, WAG, NS, and TMP wrote and edited the manuscript and figures. WAG and TMP acquired funding and supervised the project. All authors have read and approved the final manuscript.

## Availability of data and materials

All data supporting the conclusions of this article are included within the article and in additional files provided.

## Declarations

### Ethical approval and consent to participate

The patients were enrolled in protocols 76-HG-0238 and Pro00038462 approved by the National Human Genome Research Institute (NHGRI) Institutional Review Board and the Cedars-Sinai Medical Center Institutional Review Board, respectively.

## Consent for publication

Not applicable.

## Supplementary Information

See Supplementary Information Section

## Abbreviations

AAVS1: Adeno-associated virus integration site 1
CLN6: Neuronal ceroid lipofuscinosis, type 6
CLN6A: Variant late-infantile CLN6
CLN6B: CLN6 Kufs type
CLN6p: CLN6 protein
CTL: Control
CTSD: Cathepsin D
DEG: Differentially-expressed gene
DOX: Doxycycline
EB: Embryoid body
ER: Endoplasmic reticulum
FACS: Fluorescent cell activated sorting
GO: Gene Ontology
ICC: Immunocytochemistry
IPSC: Induced pluripotent stem cells
I3N-IPSCs: Integrated, inducible, and isogenic Ngn2 IPSCs
LAH: lysosomal acid hydrolase
LAMP1: lysosomal associated membrane protein-1
NCL: Neuronal ceroid lipofuscinosis
NDM: Neuronal differentiation medium
NeuN: Neuronal Nuclear Antigen
NGN2: Neurogenin-2
NPCs: Neural progenitor cells
NM: media neuronal maturation media
OPC: Oligodendrocyte Progenitor Cell
PBMCs: Peripheral blood mononuclear cells
Pre-OPC: Pre-Oligodendrocyte Progenitor Cell
PPT1: Palmitoyl-protein thioesterase-1
PDI: Protein disulfide-isomerase
PT-Day: Post-thaw day
qRT-PCR: Quantitative reverse-transcriptase-mediated polymerase chain reaction
SUBC: Subunit C of the mitochondrial ATP synthase
TPP1: Tripeptidyl peptidase-1
UPR: Unfolded Protein Response
WMSAs: White matter signal abnormalities
vLINCL: Variant late-infantile neuronal ceroid lipofuscinosis

