## Supplemental Table 1 for "Cellular Modeling of CLN6 with IPSC-derived Neurons and Glia"

**Supplementary Information**

***Clinical Information of CLN6 subjects***

Each subject underwent record review and clinical evaluation. Some subjects have been previously reported (Chin et al., 2019). Brief summaries of each subject’s genotype, phenotype and neuroimaging. Each subject had WMSAs present on brain MRI studies either early or late in their disease course (Fig 1). Family relationships are identified in parentheses below whenever present. IPSC lines were generated from some subjects and are also identified in the parentheses below. Several subjects were subsequently enrolled in an independent therapeutic clinical trial; therefore, clinical details after their enrollment were unavailable.

**Subject 1** (older sister of Subject 2 and previously reported as Proband 2) presented at 12.5 years and was identified with homozygous CLN6 variants (c.723G>T; p.M241I) by exome sequencing.(Chin et al., 2019) She had a history of normal development until age 5 years when she exhibited slowly progressive symptoms that included developmental regression, ataxia, lower limb spasticity, dystonia, and progressive myoclonic epilepsy*.*(Chin et al., 2019) Brain MRI at ~12 years showed atrophy of the cerebrum, cerebellum, basal ganglia, corpus callosum and brainstem. There was significant white matter atrophy (greater than gray matter) along with white matter signal abnormalities (WMSAs) extending into occipital lobes (Fig1).

**Subject 2** (younger brother of Subject 1) presented at 8 years with subsequent exome sequencing identifying the same homozygous CLN6 variants (c.723G>T; p.M241I). He developed normally until age 5 when he manifested a phenotypic pattern similar to that of his sister.(Chin et al., 2019) Brain MRI at ~8 years had less atrophy than his sister, but significant WMSAs were consistent with a leukodystrophy that extended into the occipital lobes (Fig1).

**Subject 3** (previously reported as Proband 1)(Chin et al., 2019) presented at 8 years and was identified with heterozygous CLN6 variants (c.218-220dupGGT; p.W73dup and c.296A>G; p.K99R) by exome sequencing. At 5 years she developed progressive myoclonic epilepsy and subsequently progressed to have ataxia and developmental regression.(Chin et al., 2019) MRI brain at ~9.5 years had significant white matter loss (compared to gray matter atrophy) associated with abnormal symmetric T2/FLAIR WMSAs that extended into her occipital lobes (Fig 1).

**Subject 4** was diagnosed by exome sequencing at 15 years with homozygous *CLN6* variants (c.768C>G; p.D256). He had met his milestones in his early years but then developed progressive ataxia, dysdiadochokinesia, dysmetria, tremor, dysarthria, and learning difficulties. He had mild lower extremity hypertonicity, spasticity and a progressive decline in fine motor skills. He never had clinical seizures although he had an abnormal EEG showing “slow and spike waves in the occipital region” and “generalized burst activity and spikes.” MRI at 11 years showed normal cerebellum and bilateral white matter signal abnormalities in the peritrigonal and deep parietal white matter. MRS was within normal limits.

**Subject 5/CLN3** (older sister of Subject 6; cellular sample: CLN3) presented at 4 years and was diagnosed by exome sequencing with heterozygous CLN6 variants (c.184C>T; p.R62C and c.486+1G>A). She met milestones until 30 months when the family noticed increased difficulty with speech and fine motor ability. She then developed seizures and myoclonus at ~4 years and subsequent EEG noted frequent bilateral occipital spike-wave discharges. These were associated with occasional paroxysmal bursts of high-amplitude, bilateral frontocentral predominant 2-2.5 Hz delta slowing. Neuro-ophthalmological evaluation, including optical coherence tomography, was normal at ~4.5 years. Neuropsychological testing at that time showed deficits in cognition, visual processing, and fine/gross motor skills, while less extensive deficits were present in her expressive and receptive language skills. MRI at ~5 years had a demyelinating leukodystrophy with symmetric confluent signal abnormality within the supratentorial white matter, ventral pons, and pyramids (Fig. 1). She was enrolled in a therapeutic clinical trial at ~6 years.

**Subject 6/CLN2** (younger sister of Subject 5; cellular sample: CLN2) was diagnosed at ~21 months with the same CLN6 variants as her sister by familial variant testing (c.184C>T; p.R62C and c.486+1G>A). She sat independently at 5-6 months and walked at 15 months. Neuropsychological testing and language development were within normal limits at 25 months. Brain MRI at ~2 years reported WMSAs in the association fibers (Fig. 1). She was enrolled in a therapeutic clinical trial at ~2.5 years.

**Subject 7** (older brother of Subject 8) was diagnosed at ~5 years with compound heterozygous variants in CLN6 (c.214G>T; p.E72X and c.486+1G>A) on an NCL-specific gene panel. He was meeting his milestones up until ~4 years when he started falling behind his peers. Early signs were inattentiveness, fine/gross motor skill loss, and myoclonus. Neuro-ophthalmological evaluation at 5 years revealed gaze-evoked nystagmus and retinal abnormalities. Neuro-ophthalmological evaluation at 5 years revealed gaze-evoked nystagmus and retinal abnormalities. Neurological progression included ataxia and gait abnormalities. MRI at ~6 years had WMSAs associated with cerebellar hemispheric and vermian volume loss (Fig. 1). He was enrolled in a therapeutic clinical trial at ~6.5 years.

**Subject 8/CLN1** (younger brother of Subject 7; cellular sample: CLN1) was diagnosed at 9 months by familial variant testing (c.214G>T; p.E72X and c.486+1G>A) secondary to his brother’s genetic diagnosis. The patient was asymptomatic and meeting his milestones at the time of diagnosis, but was identified with speech delays the following year. He was enrolled in a therapeutic clinical trial at 1 year of age; pre-enrollment neuroimaging was not available.

**Subject 9/CLN4** (cellular sample: CLN4) was diagnosed at 5 years with homozygous variants in *CLN6* (c.665+1 G>A, intron 6) using a targeted next-generation sequencing panel for epilepsy. She was a 29-week premature female with a 2.5-month NICU stay without neonatal complications. She was mildly delayed, walking at 18 months and talking “much later.” Her speech did not develop beyond a few words. She developed toe-walking and an abnormal gait. Subsequent epilepsy evaluation noted frequent seizures, especially during sleep. She had a steady neurodegenerative progression with intractable epilepsy and loss of ambulation and passed away at 9 years of age. Neuroimaging for this subject was not available.

**Supplemental Table 1.** Subject data for IPSC lines

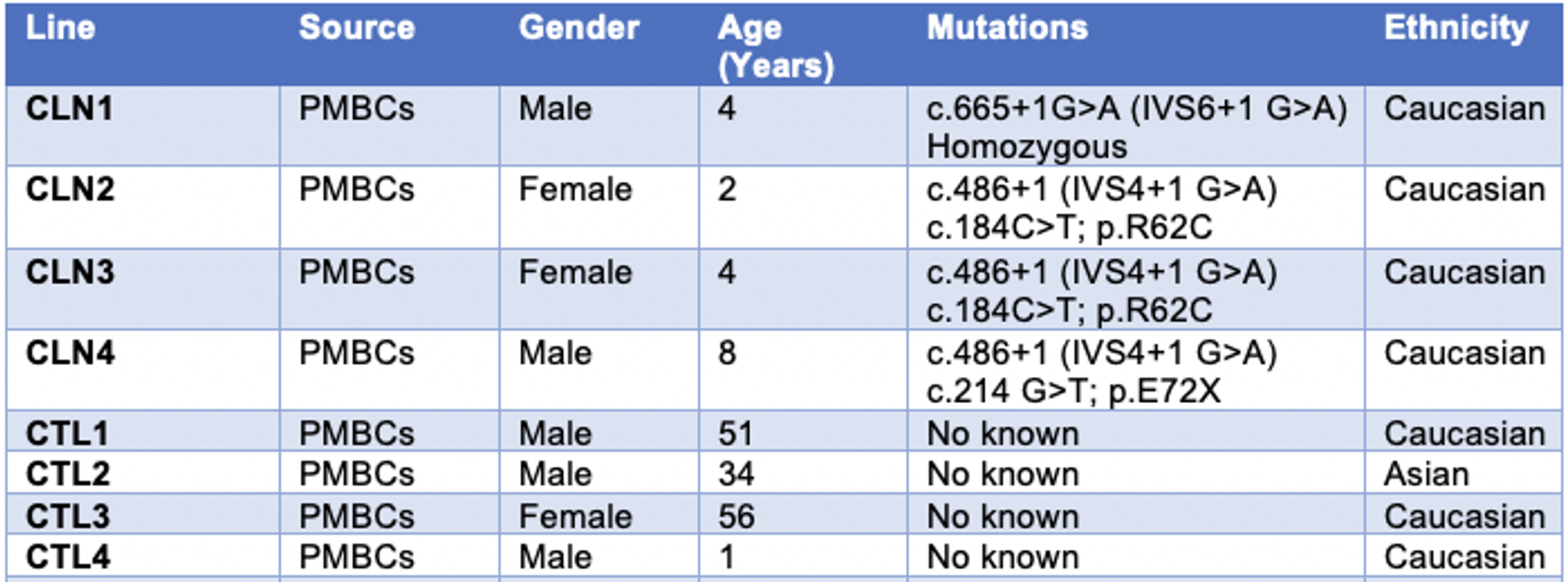

**Supplemental Table 2**

**DEGs of interest and their adjusted p-value**

|  | Gene | adj p-value |
| --- | --- | --- |
| Lysosomal Function, Size, Numbers & Formation | *LYST* | 4.1 E-07 |
|  | *SCPEP1* | 0.003 |
|  | *SMPD1* | 0.003 |
|  | *MCOLN1* | 0.015 |
|  | *CTSA* | 0.013 |
| Lysosomal-Vesicular Proton Pump Subunits | *ATP6V1H* | 0.022 |
|  | *ATP6V1G2* | 0.032 |
|  | *ATP6V0C* | 0.038 |
| Cellular Motor Subunits | *DYNC1H1* | 5.742 E-06 |
|  | *DYNC1I1* | 0.002 |
|  | *KIF3C* | 0.019 |
| Trafficking Proteins | *RAB7A* | 0.018 |
|  | *VPS39* | 0.031 |
|  | *STXBP1* | 0.041 |
| Synaptic Function | *SLC17A7* | 0.006 |
|  | *ID4* | 0.020 |
|  | *VAMP2* | 0.051 |
|  | *SNAP25* | 0.032 |
| Cytokines and Cytokine Receptors | *CX3CL1* | 0.004 |
|  | *FGFR2* | 0.024 |
|  | *EPHA7* | 0.026 |
|  | *MDK* | 0.027 |
| Neuronal Apoptosis | *C5AR1* | 0.0001 |
|  | *PINK1* | 0.004 |
|  | *TYRO3* | 0.037 |

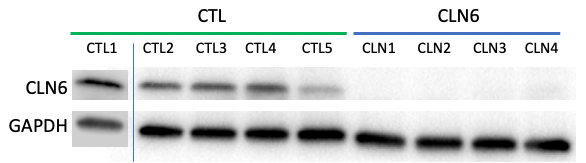

**Supplemental Figure 1.** CLN6 protein levels in IPSC lines. Western blot of CTL- and CLN6-IPSCs with anti-human CLN6p antibody (gift of Dr. Stella Lee) indicate that CLN6 protein is absent from our CLN6 lines. CTL1 results are from a separate blot. CTL5 was a heterozygote for a CLN6 variant (c.486+1 G>A). GAPDH as loading control.

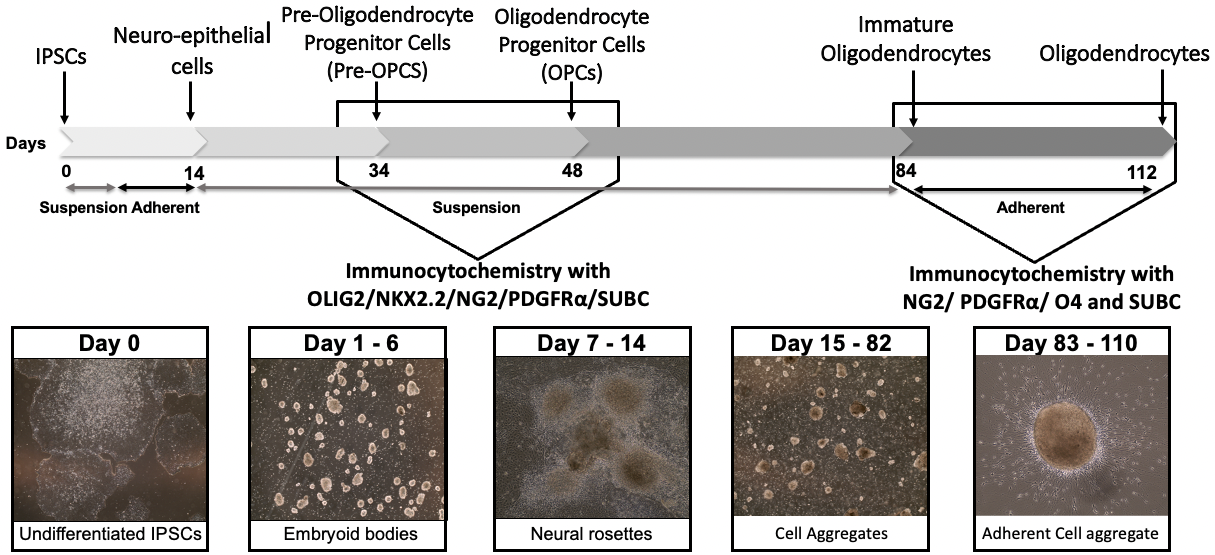

**Supplemental Figure 2. Oligodendroglial differentiation protocol.** IPSCs were grown in suspension or as adherent cells throughout the 112 days of culture. Figure indicates time-points where cells were harvested and underwent ICC for specific markers. Representative images of cultures during differentiation stages.

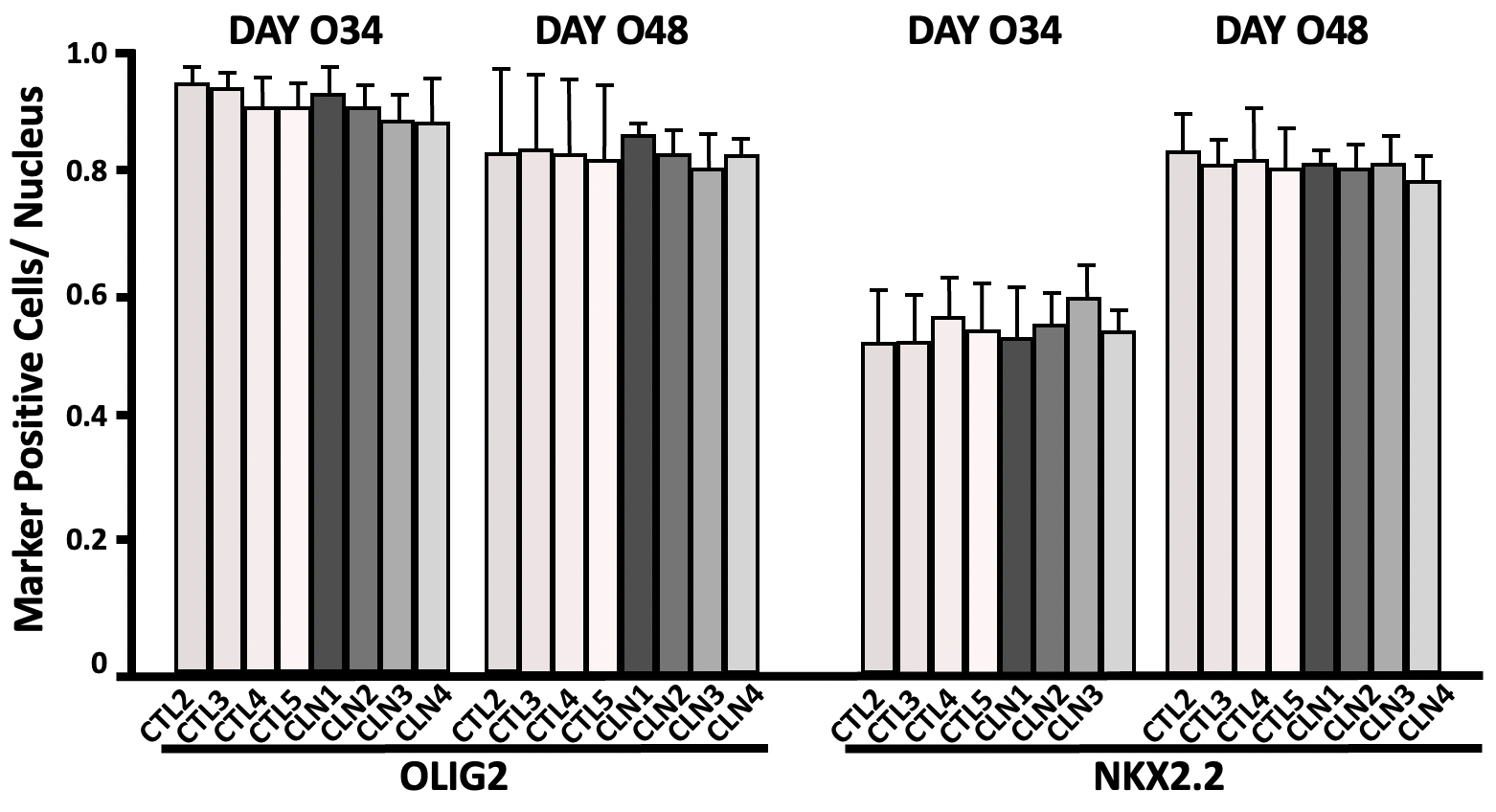

**Supplemental Figure 3.** IPSC cultures expressing oligodendroglial differentiation markers OLIG2 and NKX.2 at time-points Days O34 and O48. There were no significant differences in expression of the two markers in CTL- and CLN6 cultures at either time-point. Values represent mean ± S.D.; N = 3 biological replicates per cell line; One-Way ANOVA Bonferroni’s multiple comparison test. * p < 0.05

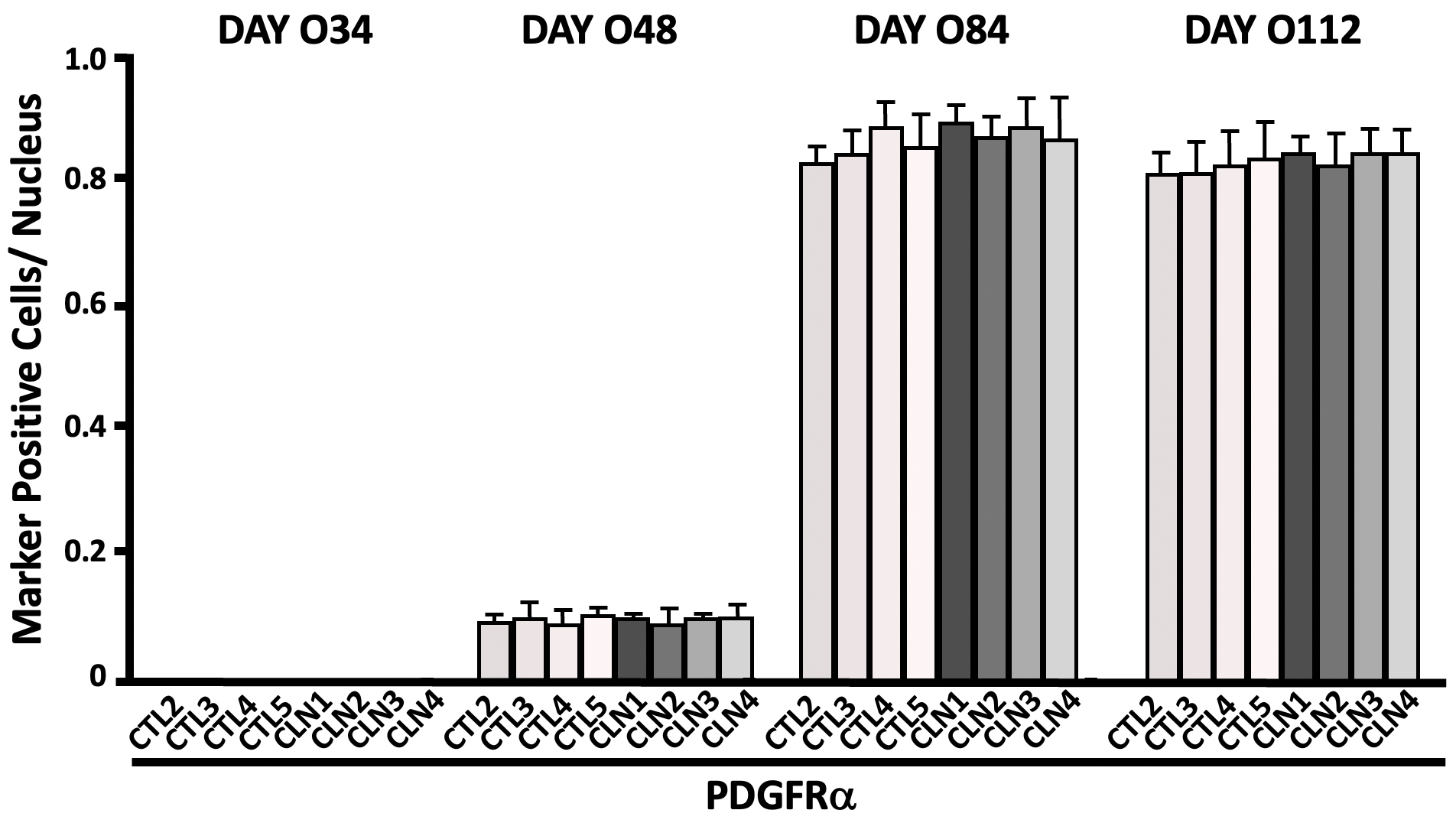

**Supplementary Figure 4.** IPSC cultures expressing oligodendroglial differentiation marker PDGFRa at time-points Days O34, O48, O84, and O112. There were no significant differences in expression of the marker in CTL- and CLN6 cultures at either time-point. Values represent mean ± S.D.; N = 3 biological replicates per cell line; One-Way ANOVA Bonferroni’s multiple comparison test * p < 0.05; ** p < 0.01; *** p < 0.001

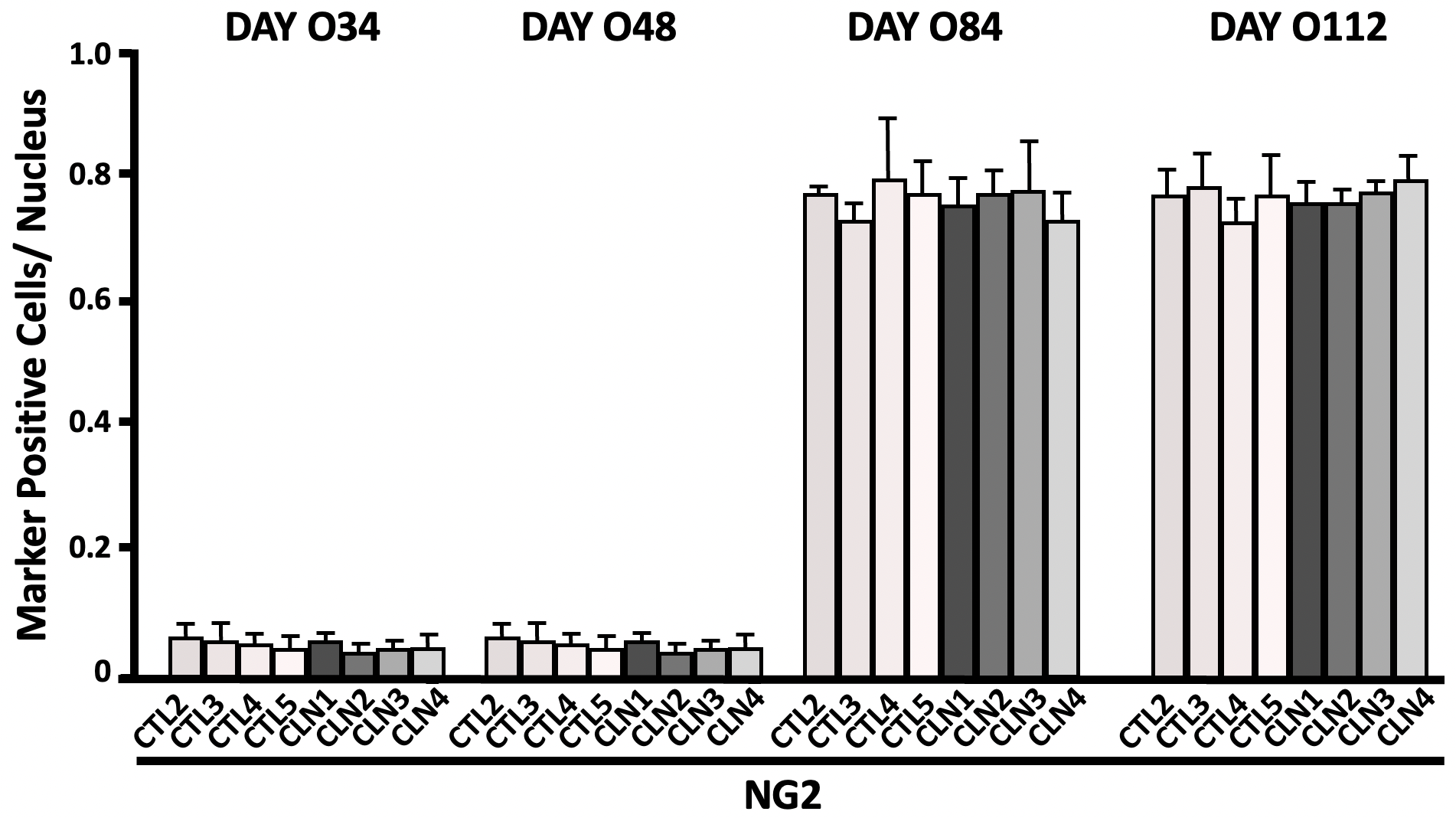

**Supplementary Figure 5.** IPSC cultures expressing oligodendroglial differentiation marker NG2 at time-points Days O34, O48, O84, and O112. There were no significant differences in expression of the marker in CTL- and CLN6 cultures at either time-point. Values represent mean ± S.D.; N = 3 biological replicates per cell line; One-Way ANOVA Bonferroni’s multiple comparison test * p < 0.05; ** p < 0.01; *** p < 0.001

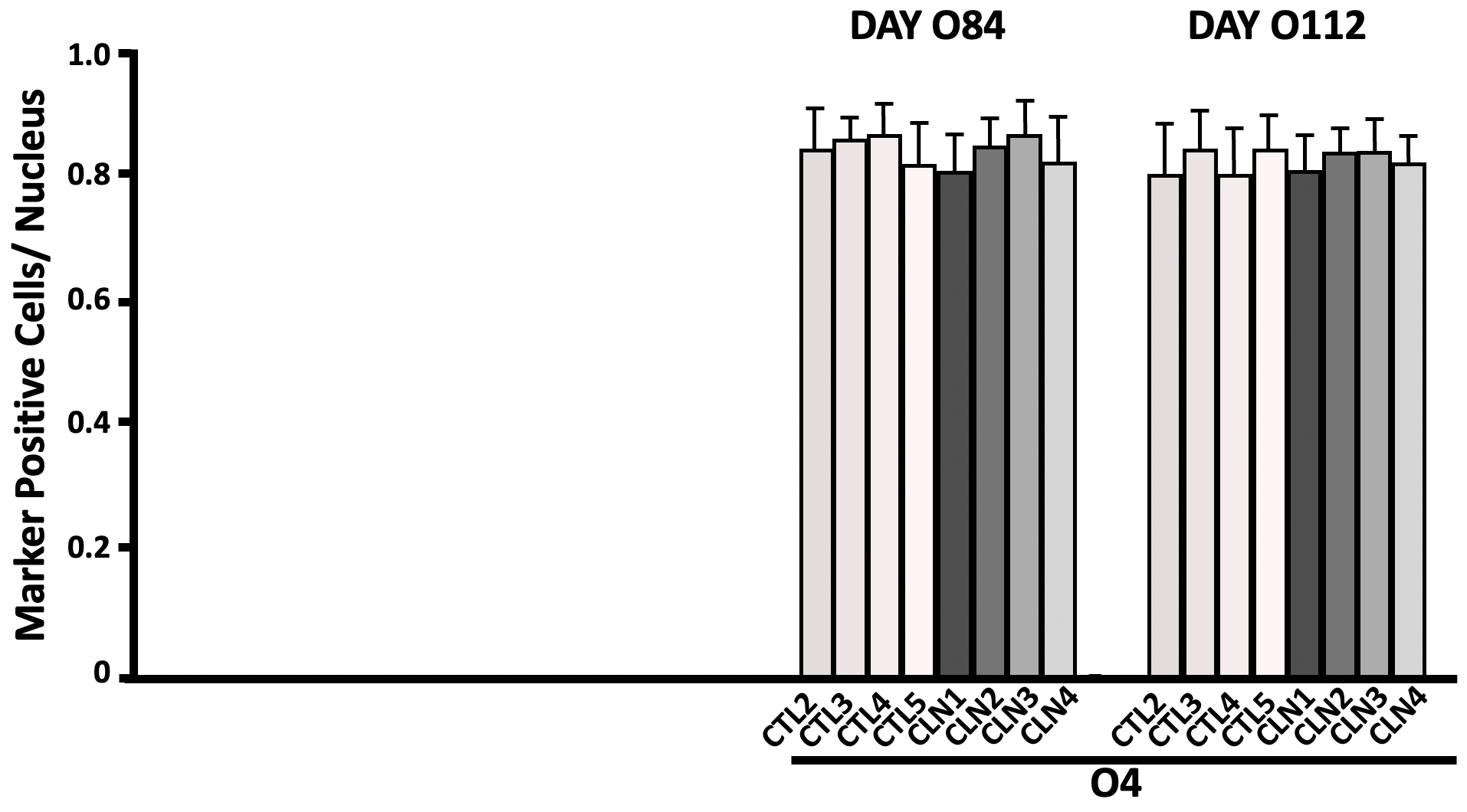

**Supplementary Figure 6.** IPSC cultures expressing oligodendroglial differentiation marker O4 at Days O84 and O112. There were no significant differences in expression of the marker in CTL- and CLN6 cultures at either time-point. Values represent mean ± S.D.; N = 3 biological replicates per cell line; One-Way ANOVA Bonferroni’s multiple comparison test * p < 0.05; ** p < 0.01; *** p < 0.001

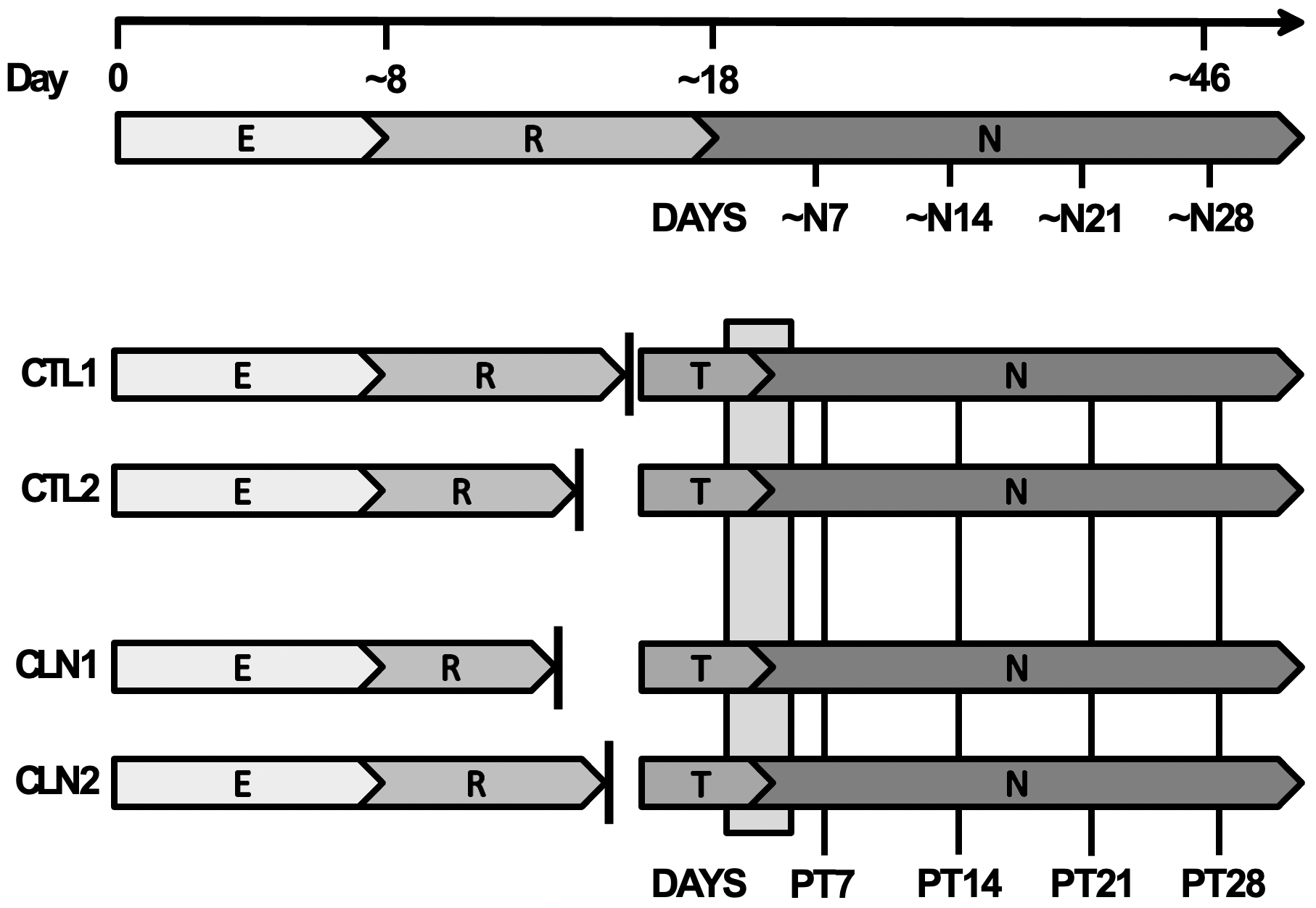

**Supplementary Figure 7. Modified neuronal differentiation protocol.** CTL- and CLN6-IPSC lines were differentiated through the epithelial (E) and neural rosette (R) stages and frozen when ~80% or more had taken on neural rosette morphological characteristics. Cells were then thawed together to “synchronize” cell lines and continued forward with neuronal terminal differentiation.

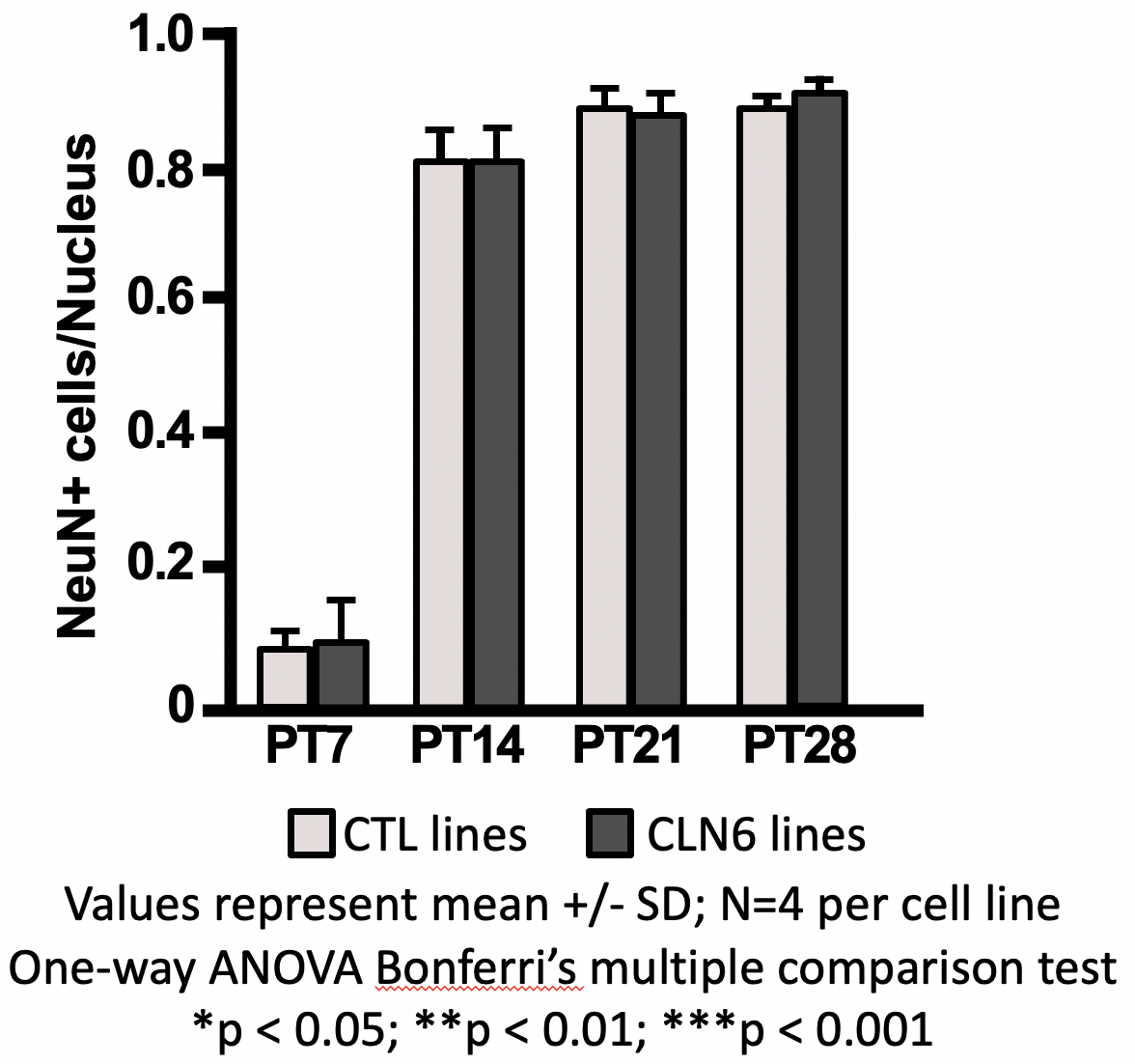

**Supplemental Figure 8.** Immunocytochemical analysis of CTL- and CLN6 cultures indicated there were no differences in the number of NeuN+ cells in these cultures.Values represent mean +/- SD; N=4 per cell line. One-way ANOVA Bonferri’s multiple comparison test. *p < 0.05; **p < 0.01; ***p < 0.001

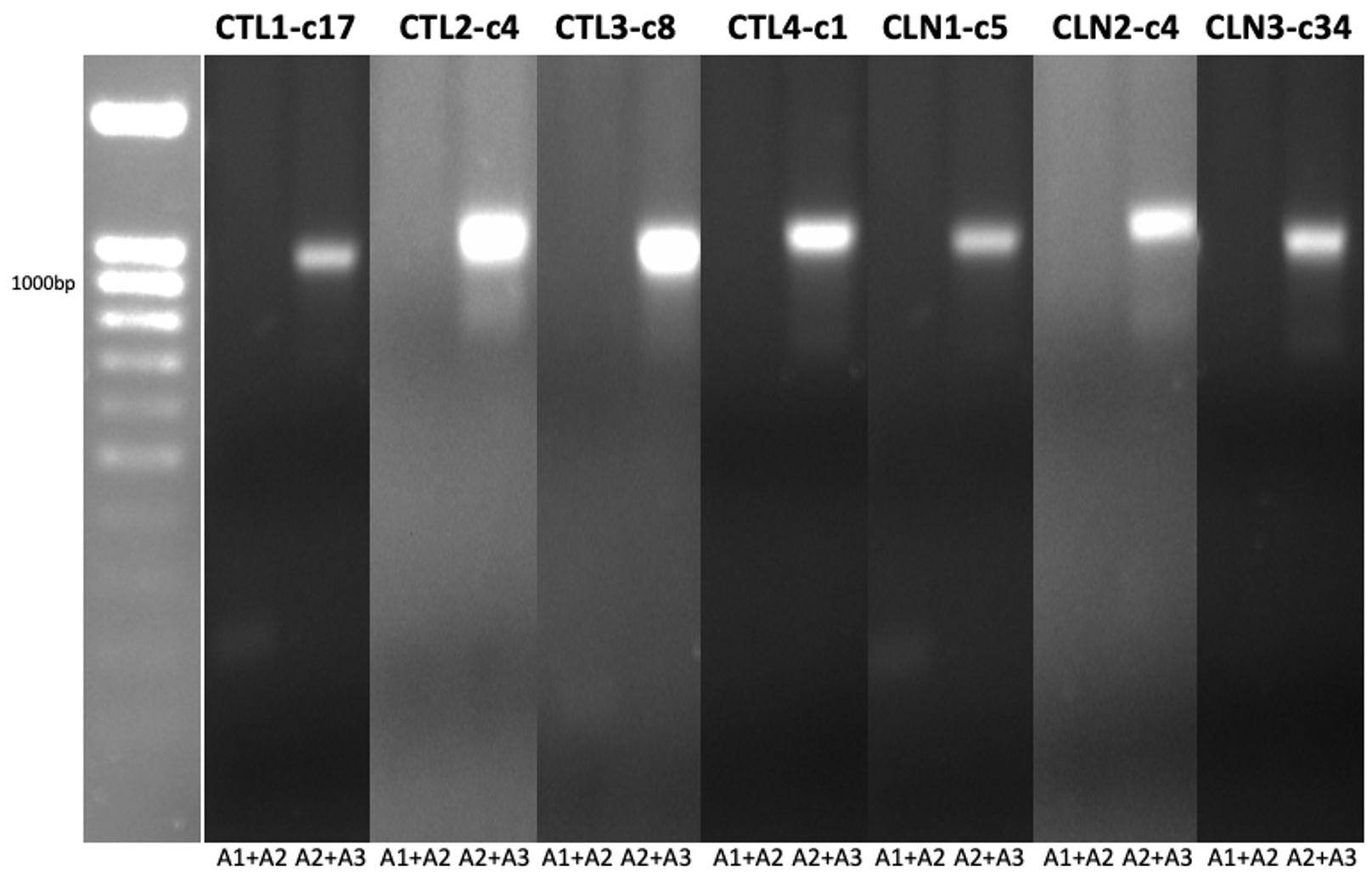

**Supplemental Figure 9. PCR analysis of CTL- and CLN6-I3N-IPSC clones.**(56) Each designated clone expressed the appropriate 1100 base-pair band with PCR primers A2+A3 for integration into the AAVS2 site. No 163 base-pair bands were seen using the A1+A2 PCR primer pairs indicating these clones are likely homozygous for the transgene insertion.

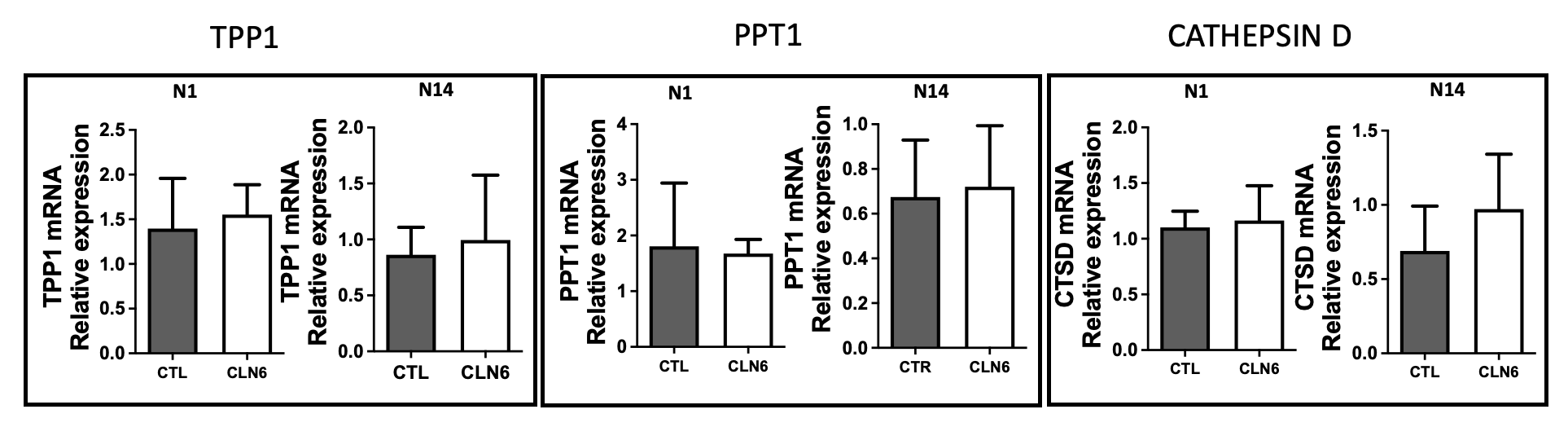

**Supplemental Figure 10. Expression of LAH transcripts.** Transcriptional evaluations of TPP1, CTSD, and palmitoyl-protein thioesterase-1 (PPT1) with quantitative reverse-transcriptase-mediated polymerase chain reaction (qRT-PCR) at Days N1 and Day N14 revealed no significant differences between CTL- and CLN6-I3N-neurons.

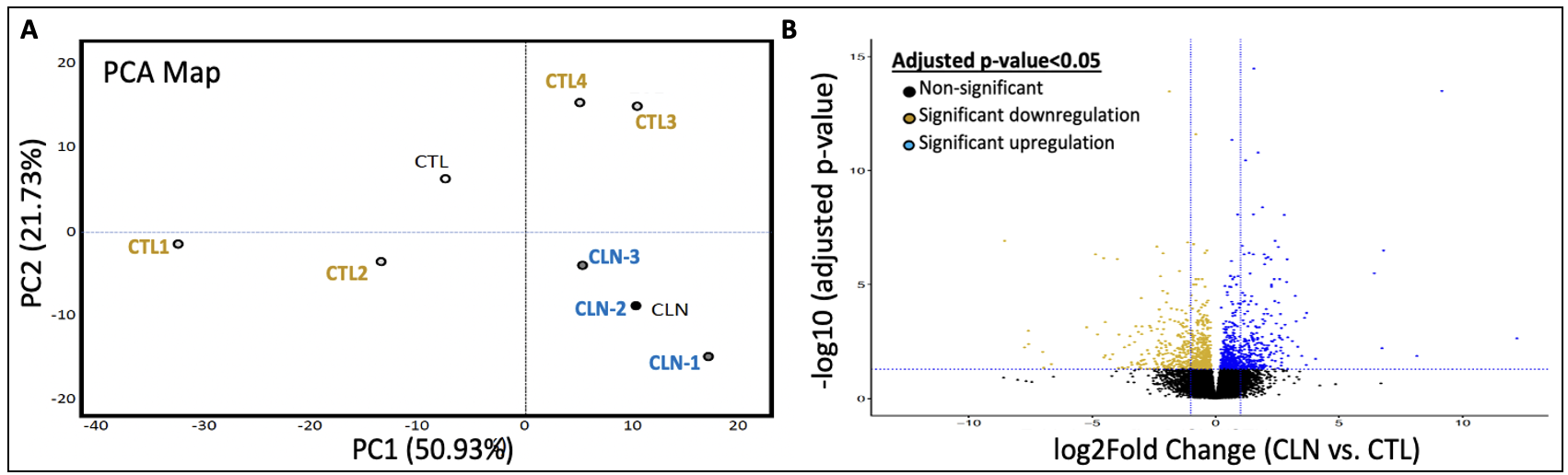

**Supplemental Figure 11. Bulk RNA-Seq in I3N-Neurons at Day N5.**
(**A**) PCA map indicates CLN6-I3N-neurons clustered together away from CTL-I3N-neurons. (**B**)Volcano plot showing 1832 total DEGs (upregulated: blue dots; downregulated: yellow dots)
